# *LCORL* and *STC2* variants increase body size and growth rate in cattle and other animals

**DOI:** 10.1101/2025.01.01.630985

**Authors:** Fengting Bai, Yudong Cai, Min Qiu, Chen Liang, Linqian Pan, Yayi Liu, Yanshuai Feng, Xuesha Cao, Qimeng Yang, Gang Ren, Shaohua Jiao, Siqi Gao, Meixuan Lu, Xihong Wang, Rasmus Heller, Johannes A. Lenstra, Yu Jiang

## Abstract

Using ancestral recombination graphs, we investigated recent selection signatures in European beef cattle breeds, pinpointing sweep-driving variants in the *LCORL* and *STC2* loci with notable effects on body size and growth rate. The ACT-to-A frameshift mutation in *LCORL* occurs mainly in central-European cattle, and stimulates growth. Remarkably, convergent truncating mutations were also found in commercial breeds of sheep, goats, pigs, horses, dogs, rabbits, and chickens. In the *STC2* gene, we identified a missense mutation (A60P) located within the conserved region across vertebrates. We validated the two natural mutations in gene-edited mouse models, where both variants in homozygous carriers increase the average weight by 11%. Our findings provide insights into a seemingly recurring gene target of body size enhancing truncating mutations across domesticated species, and offer valuable targets for gene editing-based breeding in animals.

## INTRODUCTION

European cattle body size has increased over the past 1000 years (Fig. 1B) ^1^. Body size and growth rate are still critical selection criteria in domesticated animals, particularly for meat-producing livestock. Various livestock breeds exhibit together a substantial phenotypic diversity, providing an excellent resource for exploring large-effect variants contributing to differences in body size. Because mammals share a conserved set of genes regulating body size ^2,3^, investigating domesticated animals’ body size is also relevant for human health and animal body size regulation.

**Fig. 1.**
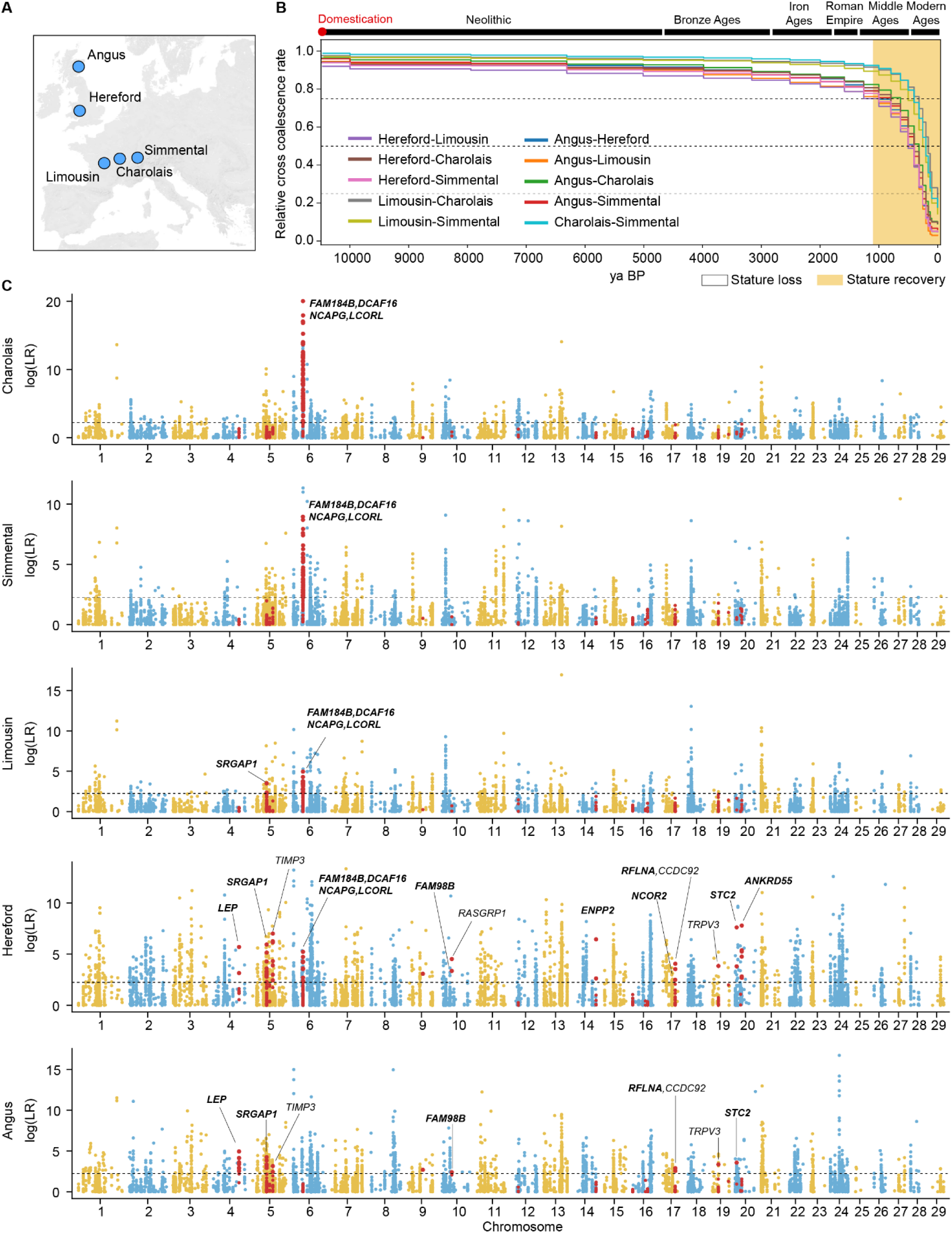
Divergence time of five European beef cattle breeds and selection analysis over the past 1,000 years. **(A)** Geographic origins of the five European beef cattle breeds studied in this research. **(B)** Relative coalescence rates among the five beef cattle breeds inferred from the ARG. Around 1,000 years ago, genetic divergence among the breeds accelerated, coinciding with a gradual recovery in the body size of domestic cattle. **(C)** CLUES analysis of highly differentiated biallelic SNVs across the five cattle breeds. Genes overlapping or nearest of the CLUES selected SNVs are annotated. Selected genes associated with human body size are in bold. The black dashed line represents the log(LR) threshold for rejecting neutral mutations. Variants located within cattle body size QTLs are highlighted in red.

Domestic taurine cattle originated from aurochs in the Fertile Crescent around 10,500 years ago^4,5^ and were introduced to Europe approximately 8,500 years ago ^6,7^. Archaeological evidence suggests that after domestication, the body size of cattle decreased until the Middle Ages, after which it gradually increased ^1^, indicating selection for larger body size, which could have left signatures of selective sweeps in the genome. Concordantly, some cattle body size and body weight QTLs identified through genome-wide analysis also exhibit signatures of selective sweeps^2^. Cattle body size traits thus present an ideal and valuable model for integrating genome-wide association studies (GWAS) with selective sweep analyses to identify sweep-driving mutations. However, previous selective sweep analyses of body size QTLs have generally focused on genomic regions rather than individual variants, limiting the resolution in localizing adaptive mutations ^2^. So far, several candidate gene variants have been identified in Australian cattle^8^, Qinchuan cattle ^9^, Wagyu ^10^, Gelbvieh ^11^, Canadian beef breeds ^12^, Red Angus ^13^ and Belgian Blue cattle ^14^. With the exception of An et al., 2019, these studies consistently localize a causative variant on bovine chromosome 6 in or near the genes *NCAPG* and *LCORL* together with several other variants. A Haplotype Encompassing two *LCORL* coding variants was proposed to be associated with increased lean growth in the Charolais breed ^15^.

In this study, we focus on five beef cattle breeds: Angus and Hereford from Britain, and Charolais, Limousin, and Simmental from Continental Europe. These breeds exhibit systematic differences in body size and weight, with British cattle generally being smaller and lighter compared to their Continental European cattle ^16,17^. Utilizing the advanced ancestral recombination graphs (ARG) ^18^ and the CLUES analyses ^19^, which facilitates selection scans at the single nucleotide variant (SNV) level, we screen for highly differentiated variants between Britain and Continental Europe cattle to assess evidence of selection within the last millennium. We identify several variants likely undergoing selective sweeps and contributing to population differentiation in body size between British and Continental beef cattle. Subsequently, we focus on two gene loci that exhibited the highest significance of recent selection and a substantial impact on growth rate. By integrating trans-ancestral selective sweep analysis, we pinpoint the variants most likely driving selective sweeps in these regions. Two new coding variants in the genes *LCORL* and *STC2* were functionally validated in gene-edited mice. For *LCORL,* convergent selection was indicated by similar variants in seven other domestic species.

## RESULTS

### Analysis of recent selection based on ancestral recombination graphs

In this study, we constructed ARG using SNVs and estimated relative cross-coalescence rates among five European beef cattle breeds: Angus, Hereford, Limousin, Simmental, and Charolais (Fig. 1A). The relative cross-coalescence rate suggested a decline to 0.5 between British cattle and Continental European cattle at 631 years ago (0.25 to 0.75 range = 200 to 1259 years ago) (Fig. 1B). This suggests that recent strong artificial selection has driven rapid genetic divergence among these breeds. We used *Fst* statistics to assess the degree of differentiation for each biallelic SNV between British and central-continental breeds (Fig. S1), defining the top 0.05% of SNVs (12,753 in total) as highly differentiated SNVs. We used CLUES to reconstruct frequency trajectories of SNVs across the five breeds and calculated the likelihood ratio (logLR) for selection versus neutrality over the last 1000 years (Fig. 1C). To reduce false selection signals due to genetic drift, we simulated the demographic history of the five beef cattle and established the logLR threshold for rejecting neutrality (cutoff=2.23) (Fig. S2). This allowed the selection of 3,208 SNVs from 12,753 highly differentiated SNVs located in 354 loci (Fig. 1C, Table S1).

In order to correlate these SNVs with body size traits, either as causal variants or in strong linkage disequilibrium with causal variants, we compared their locations with those of 158 known cattle body size QTLs^2^ (Table S2). We found 111 CLUES-identified selective SNVs located within 11 body size QTLs (Fig. 1C, Table S3). Of the 18 genes overlapping or nearest of the 111 CLUES-identified selective SNVs, 12 were found to be associated with body height in humans ^20,21^, more than expected by chance (*P* < 4 × 10^−9^, chi-squared test). The most significant associations were found for the body-size QTL regions containing the *NCAPG-LCORL* and *STC2* loci. Notably, in a previously reported GWAS of average daily gain in a mixed population comprising these five cattle ancestries ^12^, the *NCAPG-LCORL* and *STC2* loci were identified as the first and third most significant loci, respectively, with effect sizes of 71 grams/day and 31 grams/day. This suggests large effects on both body size and growth rate. Consequently, we conducted further in-depth analyses of the *NCAPG-LCORL* and *STC2* loci to investigate the variants driving their selective sweeps and population differentiation.

### Identification of the sweep-driving mutation at the *NCAPG-LCORL* locus

The *NCAPG-LCORL* region, located in cattle body size QTL Chr6: 37,180,233-37,768,454, was identified as the most significant region in our CLUES analysis for Charolais and Simmental cattle (Fig. 1C), with the selected mutations showing high frequency (frequency >= 0.69) (Fig. 2A, Table S3). However, in Angus and Hereford cattle from Britain, there is either no significant selection signal or the frequency of the selected mutation is very low (frequency <= 0.1) (Fig. 2A, Table S3). Within the Chr6: 37,180,233-37,768,454 region, 85 SNVs exceeded the genome-wide *Fst* threshold (top 0.05%), 65 SNVs of which displayed CLUES-identified selection signals that were prioritized for further analysis. We hypothesize that the sweep-driving allele in this region has a single common origin across different breeds. The spread of this allele to various breeds has resulted in heterogeneous LD patterns with the sweep-driving allele. Consequently, the true sweep-driving allele should exhibit selection signals in all four breeds that show evidence of selective sweep in the *NCAPG-LCORL* region. In contrast, alleles that have been hitchhiked under genetic drift may only show selection signals in certain breeds. Therefore, trans-ancestral analyses that combine CLUES results from different breeds can aid fine-mapping by capitalizing on ancestral differences in LD patterns. Among CLUES-identified selected SNVs, five SNVs spanning 89 kb (Chr6: 37,349,373-37,438,248) shared CLUES selection signals among these four breeds that show selective sweep in the *NCAPG-LCORL* region (Fig. 2A, Table S3). These five shared selected mutations define a narrower selection-targeted haplotype. Across the four breeds exhibiting selective sweep signals at the *NCAPG-LCORL* locus, 315 haplotypes (D haplotypes) carry the derived alleles of these five shared mutations, while 425 haplotypes (A haplotypes) carry the ancestral alleles (Fig. 2B). The D haplotypes are mainly found in Charolais and Simmental cattle, but not in the Angus (Fig. S3). Between the A haplotypes and D haplotypes, in addition to the five shared selected SNVs, three INDELs also exhibited the top frequency differences (Δ Frequency=0.98-0.99) (Fig. 2C, Table S4). All the five shared selected SNVs and two of the three top differentiated INDELs were located in non-coding regions, and the left one is a frameshift INDEL (rs384548488, ACT-to-A) (Table S5). The 2 bp mutation on rs384548488, is located in the “PRC2-associated LCORL isoform 2” (PALI2, encoded by an alternative transcript of *LCORL*), resulting in the complete loss of the PIP (PALI interaction with PRC2) domain in PALI2 (Fig. 2D), which is expected to interfere with PRC2 (Polycomb Repressive Complex 2) methyltransferase activity ^22^. Using the GWAS results for body weight in Red Angus and average daily gain in a mixed cattle population ^12,13^, we inferred that the level of LD (r²) with rs384548488 correlates strongly (r > 0.76, *p* < 1 x 10^-25^) with the-log_10_(GWAS *p-value*) of the variants (Fig. 2E, Fig. S4). This suggests that rs384548488 is strongly associated with growth trait variations in several beef populations (see below for its geographic range). Therefore, rs384548488*A, a predicted loss-of-PIP-domain (pLoPD) mutation, is highly likely the sweep-driving allele and the causal variant affecting cattle body size and growth rate at the *NCAPG-LCORL* locus.

**Fig. 2.**
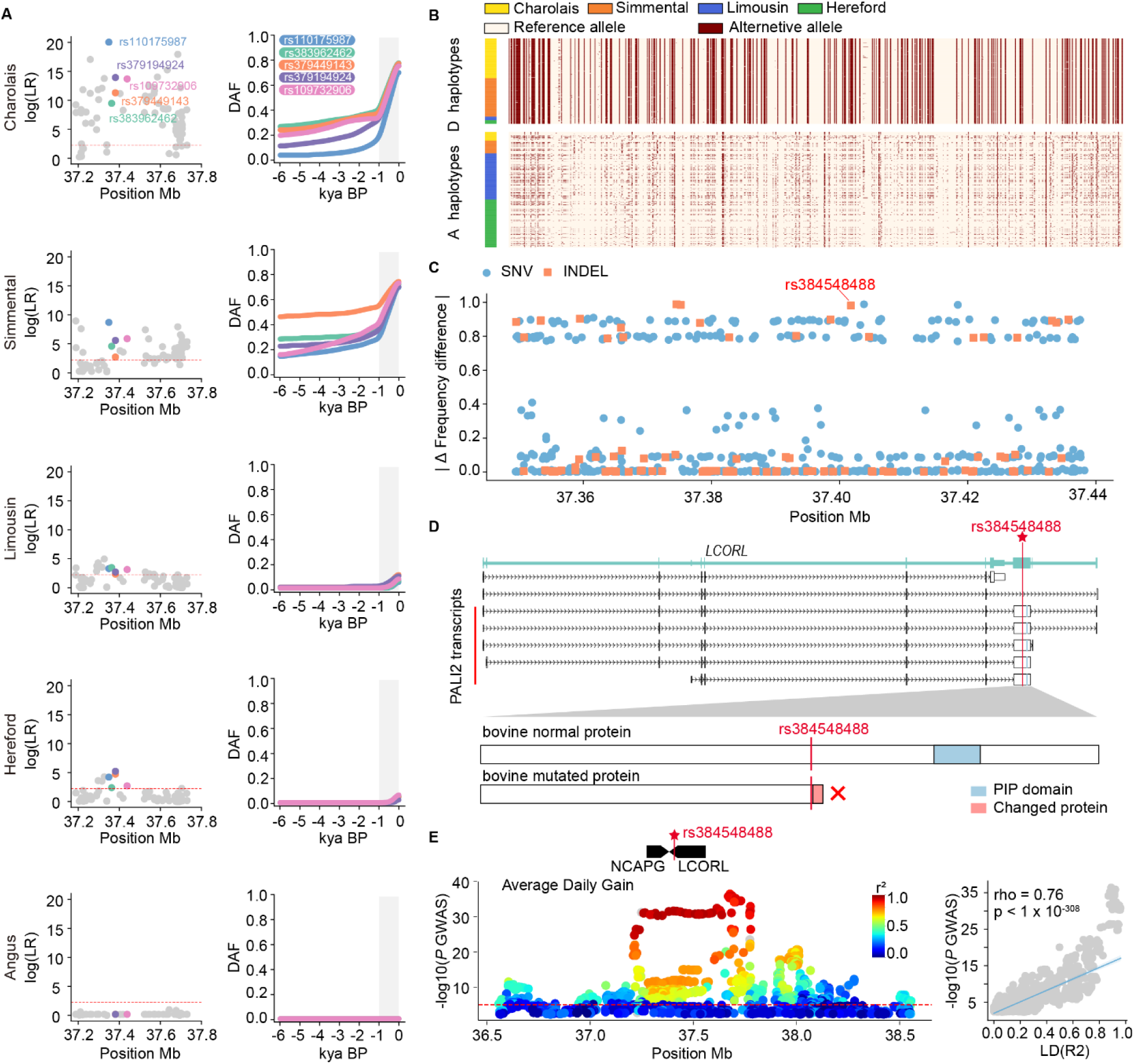
The rs384548488 mutation at *LCORL* is a candidate variant for cattle growth in the *NCAPG-LCORL* region. **(A)** Detailed plots for Chr6:37180233-37817575 encompassing the *NCAPG-LCORL* locus. The first column presents zoomed-in Manhattan plots of log(LR) for each breed. The second column shows allele frequency trajectories for the five shared selective SNVs in the CLUES analysis. Gray shading highlights the time range from 1,000 years ago to the present. **(B)** A clear difference is observed between D and A haplotypes, and the notably lower diversity of D haplotypes suggests a selective sweep. The alternative allele relative to the reference genome is indicated in red. D haplotypes (n=315), A haplotypes (n=425). **(C)** The absolute frequency difference between D and A haplotypes. SNVs are represented by circles, while INDELs are represented by squares. **(D)** Schematic of the *LCORL* gene structure and its transcripts. The frameshift INDEL (rs384548488) mutation is highlighted by a red vertical line. Below the gene structure is a schematic of the truncation in the PALI2 protein, an isoform expression product of *LCORL*, caused by rs384548488. **(E)** The rs384548488 variant is associated with cattle average daily gain variation. The left shows the *NCAPG-LCORL* locus as a QTL for average daily gain^12^. The red line is the nominal significance at *p* = 1 x 10^-5^. The right demonstrates a significant positive correlation between the-log_10_(*p-value*) of variants in average daily gain GWAS and their LD (R²) with rs384548488, indicating that rs384548488 is linked to growth traits.

### A pLoPD mutation similar to rs384548488*A increases body size and weight in mice

To test this hypothesis, we used CRISPR-Cas9 to introduce a pLoPD mutation in C57BL/6J mice, mirroring mutation rs384548488 in cattle (Fig. S5). We weighed wild-type (+/+), heterozygous (+/-), and homozygous (-/-) PALI2 PIP domain knockout mice from weaning to 9 weeks of age (Fig. 3A). The PALI2^-/-^ mice are significantly heavier than PALI2^+/+^ mice. The effect of the pLoPD mutation on body weight displayed a dosage-dependent trend. At nine weeks of age, PALI2^+/-^ and PALI2^-/-^ males were respectively 5.9% (1.5 grams) and 9.4% (2.4 grams) heavier than PALI2^+/+^ males. Similarly, PALI2^+/-^ and PALI2^-/-^ females were respectively heavier 4.8% (1.0 grams) and 12.4% (2.6 grams) heavier than PALI2^+/+^ females. At 7 weeks of age, the body length of PALI2^-/-^ males and females was also significantly increased, with a 3.0% (2.9 mm) and 2.6% (2.3 mm) increase, respectively, compared to wild-type mice (Fig. 3B). We next analyzed mouse embryos at the E14 stage, as well as on muscle, and thymus tissues from adult mice by western blot, to determine whether increased growth rates are related to changes in H3K27me3 levels (Fig. 3C). The H3K27me3 levels were found to be reduced in PALI2^-/-^ mice compared to wild-type mice. This result confirms the possible function of PALI2 as a crucial component of the PRC2 complex ^22^, which mediated H3K27me3 in regulating individual development and growth^23–25^.

**Fig. 3.**
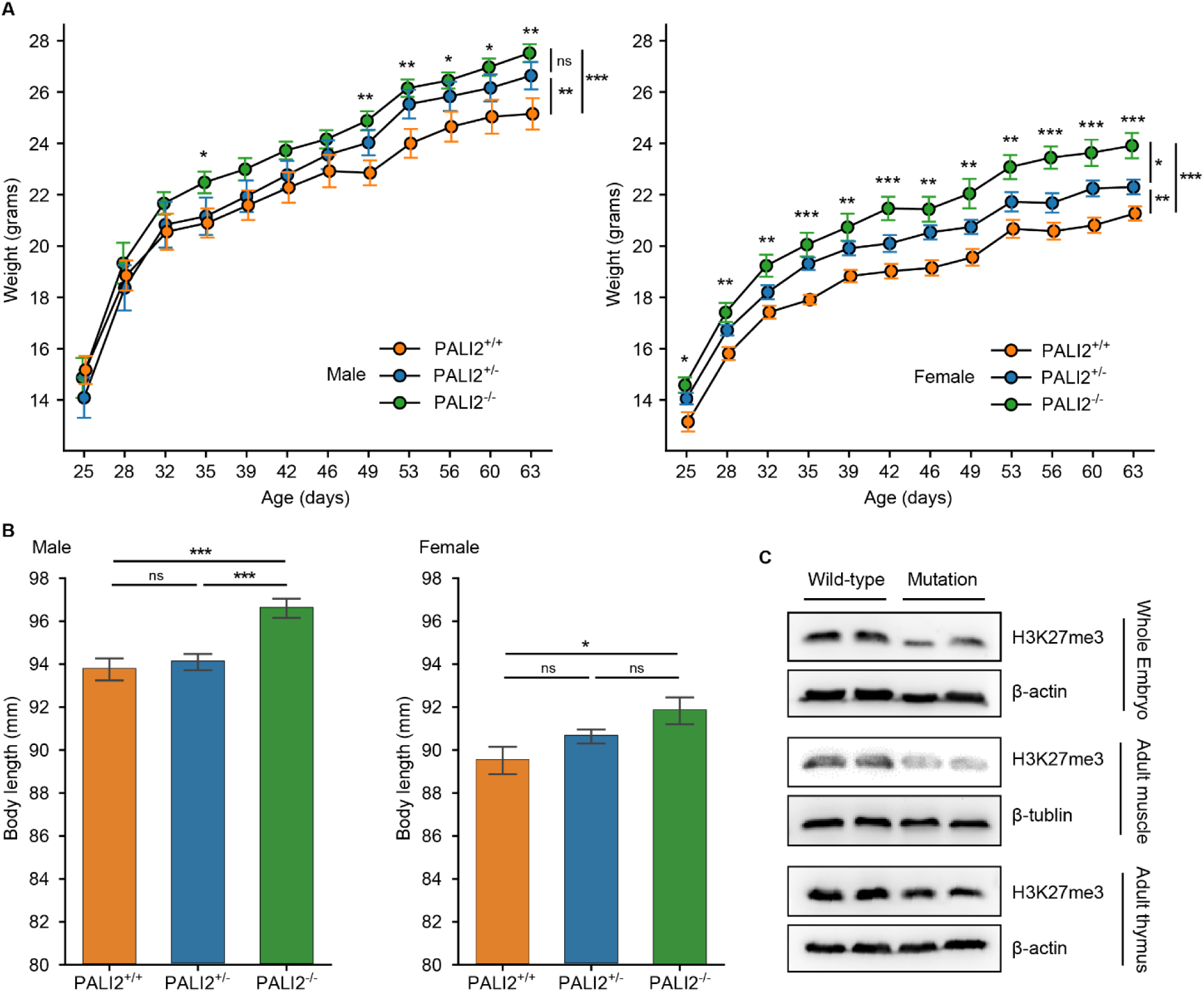
The pLoPD mutation increase body weight and body length in mice. (A) Total body mass of wild-type (+/+), heterozygous (+/-), and homozygous (-/-) PALI2 PIP domain knockout mice from 3 to 9 weeks of age. Male n = 10, 10, 10; Female n = 10, 9, 9. Two-sided t-tests were used to infer significant body mass differences between PALI2^-/-^ and PALI2^+/+^ mice at each time point (notations above each time point). Subsequently, for male mice (49 to 63 days) and female mice (25 to 63 days), repeated measures two-way ANOVA followed by Tukey’s post hoc multiple comparison test was conducted to infer significant body mass differences among the three genotypes (notations on the far right). (B) Body length of PALI2^+/+^, PALI2^+/-^, and PALI2^-/-^ mice at 7 weeks of age. Male n = 14, 23, 19; Female n = 8, 32, 9. Error bars represent mean ± standard error; one-way ANOVA followed by Tukey’s post hoc multiple comparison test was used. (C) Western blots of whole-cell lysates from mouse embryos at the E14 stage, as well as muscle and thymus tissues from adult mice, probed with the indicated antibodies. Error bars represent mean ± standard error; \**p* < 0.05; \*\**p* < 0.005; \*\*\**p* < 0.0005.

### Convergent artificial selection of pLoPD mutation in eight domesticated animals

Interestingly, we observed that several other domestic animals exhibit the loss of the PIP domain in PALI2 (Fig. 4). Reference genomes of domestic pigs (Sscrofa11.1), sheep (ARS-UI_Ramb_v2.0), and rabbits (OryCun2.0) all have a pLoPD mutation upstream of the PIP domain in PALI2 (Fig. S6, S7). Comparing domestic and wild populations suggest a selective sweep in *LCORL* locus in European domestic pigs and rabbits ^26,27^. However, these studies have not pinpointed the sweep-driving variants within the *LCORL* locus.

**Fig. 4.**
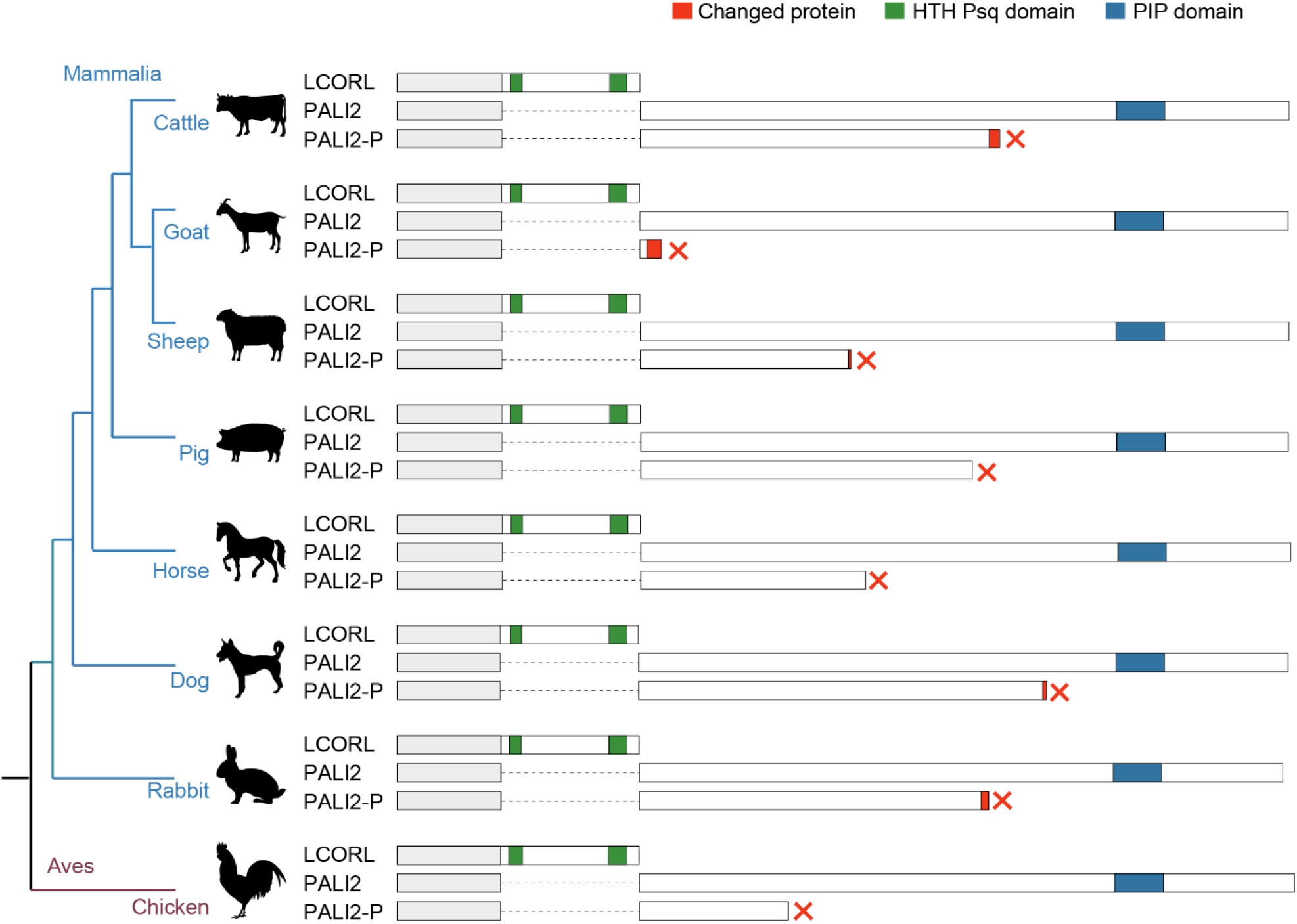
The loss of the PIP domain in PALI2 transcript across 8 domesticated animals. Natural mutations leading to PALI2 protein truncation are observed in cattle, goats, sheep, pigs, horses, dogs, rabbits, and chickens, without affecting the LCORL protein (see Fig. S4A and S5). PALI2-P represents the premature PALI2 protein. The HTH Psq domain unique to the LCORL protein is highlighted in green, while the PIP domain, specific to the PALI2 protein, is shown in blue. The regions altered by truncating mutations are marked in red, and the parts shared by all forms are colored in grey.

In pigs, the pLoPD mutation is Chr8:g.12,829,718T[6] (ancestral allele: Chr8:g.12,829,718T[7], rs697231176). The pLoPD mutation also exhibited differentiation between wild boars and European domestic pigs (Fig. S8, Table S6). The pLoPD allele at rs697231176 is found only in one Ukrainian wild boar but is absent in all 43 other European and Asian wild boars (Fig. S8A). In domestic pigs, the pLoPD allele at rs697231176 is nearly fixed in major European meat breeds (Fig. S8B).

In rabbits, the pLoPD mutation is Chr2:g.8433339 (ancestral allele:Chr2:g.8433339del). The frequency of the pLoPD allele is higher in domestic European rabbits compared to wild European rabbits, with the exception of the domestic Dutch rabbit (Fig. S9, Table S7). In the Dutch rabbit, the frequency of the pLoPD allele at Chr2:g.8433339 is only 0.11, which correlates with the smaller body size observed in this breed (Fig. S9).

In sheep, the pLoPD mutation is Chr6:g.38076935_38076936 (ancestral allele: Chr6:g.38076935_38076936insCCTGGTGGTA). The pLoPD allele is absent in wild sheep (Fig. S10A, Table S8). Moreover, extended haplotypes carrying the pLoPD allele are longer than those with the ancestral allele in Merino sheep (Fig. S10B).

We observed the same occurrences in domesticated chickens (Fig. S11). Reference genomes GRCg7b and GRCg6a lack annotations for LCORL isoforms containing the PIP domain.

However, PALI2 has been successfully annotated in reference genomes GRCg7w (Fig. S11). We hypothesize that pLoPD variants in reference genomes, has led to incorrect annotations.

According to the pangenome representing the genomes of 30 chickens^28^, a nonsense variant rs317817652 (NC_052576.1:g.75386062C>T, XP_046772233.1:p.Arg626Ter) results in the complete loss of the PIP domain in chicken PALI2 (Fig. S11). Confirming our hypothesis, reference genomes GRCg7b and GRCg6a both carry the rs317817652*T nonsense mutation. The pLoPD mutation rs317817652*T is also common in the wild species *Gallus gallus spadiceus* (frequency=0.19), from which domesticated chickens are derived (Table S9). However, this variant is absent in the other four subspecies of red jungle fowl. A GWAS on body weight and size in Chinese local chickens^29^ show that rs317817652*T significantly increases body weight and size (Table S10).

We also found pLoPD mutations in goats, horses and dogs (Fig. 4, Table S11). Recent studies reported that pLoPD mutations are associated with increased body size ^3,30,31^. The loss of the PIP domain in PALI2 in domestic cattle, goats, sheep, horses, pigs, dogs, rabbits, and chickens suggests a convergent artificial selection of a large body size and fast growth rates in domestic animals.

### Two candidate variants underlie genetic associations in the *STC2* locus

*STC2* is located within body size QTL Chr20:4967596-5008911. Within this region, the intergenic variant rs110540352 and the missense variant rs42661323*G have the highest *F*_ST_ values (Fig. S12A) and display CLUES selection signals in Hereford and Angus (Fig. 5A) In Hereford cattle, both mutations are in high linkage disequilibrium (R2 = 0.92): 96.7% of Hereford haplotypes with the derived allele rs110540352*G also carry the derived allele rs42661323*G (Fig. 5B).

**Fig. 5.**
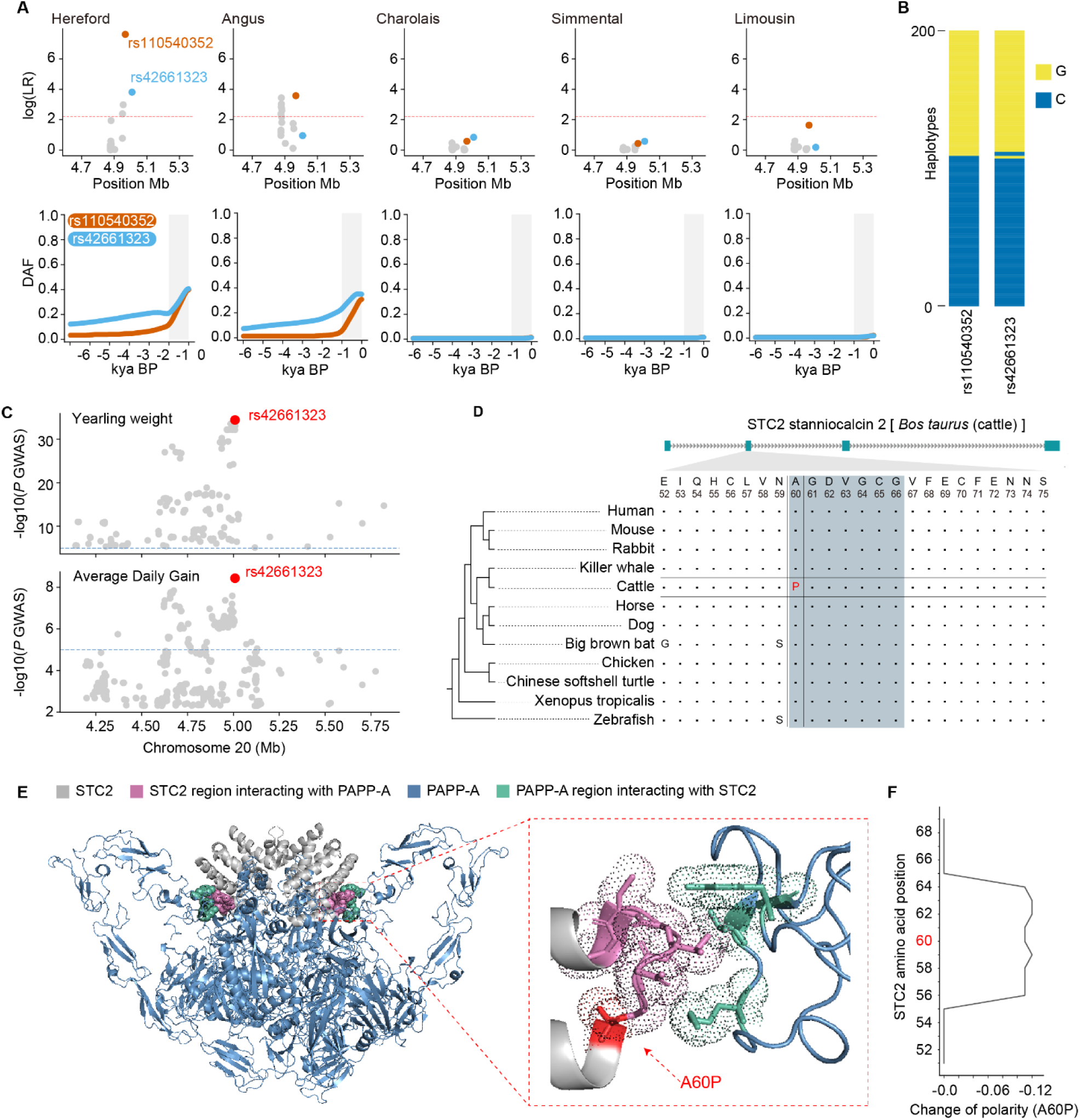
Selective sweep and functional analysis of *STC2* variants. **(A)** Detailed plots for Chr20:4598655-5366902, containing the *STC2* locus. The first row shows zoomed Manhattan plots of the log(LR) for each breed. The second row shows allele frequency trajectories for the top SNVs in CLUES analysis and missense SNVs. The gray shading indicates the time range from 1,000 years ago to the present. **(B)** Stacked bar plot of the haplotypes comprising rs110540352 and rs42661323 in the Hereford cattle (n = 200 haplotypes). The two derived alleles (rs110540352*G, rs42661323*G) are almost always inherited together as a single haplotype. The x-axis represents the number of haplotypes, and the y-axis represents the indicated SNPs. **(C)** rs42661323 is the lead GWAS SNP for yearling weight and average daily gain in cattle. The GWAS data source is the same as in Fig. 2C and Fig. S3. **(D)** Conservation analysis of the amino acid residue at position 60 of STC2. The amino acid residues at positions 60-66 of STC2 interacts with PAPP-A, marked with blue shading. **(E)** STC2 A60P may affect van der Waals interactions between STC2 and PAPP-A. Left figure: 3D structure of the 2:2 PAPP-A·STC2 complex. Middle figure: STC2 amino acids 60-66 interact through van der Waals forces with the hydrophobic pocket formed by PAPPA residues Y1566, T1594, and K1592. **(F)** Polarity prediction of STC2. Amino acid substitution Ala to Pro at position 60 of STC2 reduces the polarity from amino acids 56 to 64.

In a GWAS for cattle body size, rs110540352 43,529 bp upstream of the STC2 transcription start site (Fig. S13A) is the lead SNV at body size QTL Chr20:4967596-5008911, and the derived allele rs110540352*G shows a positive effect (β=+0.16 cm) on body size ^2^ (Table S2). We found that rs110540352 is the CLUES lead SNV in Hereford and Angus cattle, which was also indicated by CLUES selection tests on all SNVs (Fig. S12B). In mice and humans, *STC2* reduces bioactivity of insulin-like growth factors (IGFs) and inhibits growth by binding to and inhibiting PAPP-A, which cleaves IGFBPs ^32,33^. Therefore, we hypothesize that the C-to-G mutation at rs110540352 decreases *STC2* expression. According to functional annotations of the cattle genome in FAANG, rs110540352 is located in an active regulatory element (Chr20:4967400- 4969800) in the cerebral cortex and hypothalamus ^34^ (Fig. S12A). Additionally, the C-to-G mutation at rs110540352 disrupts the binding motif of E2F4 (Fig. S12B). Dual-luciferase reporter assays show that the C to G mutation at rs110540352 reduces luciferase expression (Fig. S13C). For Hereford cattle heterozygous for the rs42661323 missense variant. RNA-seq of the cerebral cortex showed that the derived allele rs42661323*G (n=7) has a significantly lower expression level than the ancestral C allele (n=16) (Binomial Test, *p*=0.047) (Fig. S13D). The high linkage disequilibrium between rs42661323*G and rs110540352*G in Hereford implies that rs110540352 has allele-specific expression, which supports that its G allele reduces *STC2* expression.

The missense C-to-G mutation at rs42661323 causes an amino acid change from alanine to proline at position 60 in STC2. In GWAS for cattle yearling weight and average daily gain, rs42661323 is the lead SNV in body size QTL Chr20:4967596-5008911 ^12,13^ (Fig. 5C). The mutation rs42661323*G increases yearling weight (β=+18.10 kilogram) and average daily gain (β=+31.34 gram). STC2 A60 is highly conserved in vertebrates except the coelacanth and the yellowbelly pufferfish (Fig 5D, Fig. S13). Structural analysis of the human PAPP-A·STC2 complex ^35^ indicates that amino acids 60-66 in STC2 interact with PAPP-A (Fig. 5E), with STC2 V63 engaging a hydrophobic pocket in PAPP-A through van der Waals forces. STC2 A60P located in this interaction region, is predicted to alter the polarity of amino acids 60-66 (Fig. 5F). Therefore, we speculate that STC2 A60P reduces STC2 binding to PAPP-A, increasing IGF bioactivity and promoting growth. This is confirmed by the prediction of program BeAtMuSiC that STC2 A60P decreases the binding affinity between STC2 and PAPP-A (ΔΔGBind=0.65 kcal/mol).

### STC2 A60P increases body size and weight in mice

To validate the effect of the STC2 A60P mutation on body size *in vivo*, we introduced this point mutation into C57BL/6J mice using the CRISPR-Cas9 system (Fig. S15). We monitored the body weights of wild-type (*STC2*^+/+^), heterozygous (*STC2*^A60P/+^), and homozygous (*STC2*^A60P/A60P^) mice from weaning until 9 weeks of age (Fig. 6C). *STC2*^A60P/A60P^ mice are significantly heavier than *STC2* ^+/+^ mice starting at 3 weeks of age (male: *p*=0.002; female: *p*=0.0001, repeated measures two-way ANOVA followed by Tukey’s post hoc multiple comparison test). The impact of the STC2 A60P mutation on body weight showed a dose-dependent trend. At nine weeks of age, *STC2*^A60P/+^ males and females were respectively 6.5% (1.8 grams) and 3.6% (0.7 grams) heavier than *STC2*^+/+^ mice, while *STC2*^A60P/A60P^ males and females were respectively 10.5% (2.8 grams) and 11.4% (2.3 grams) heavier than *STC2*^A60P/A60P^.

**Fig. 6.**
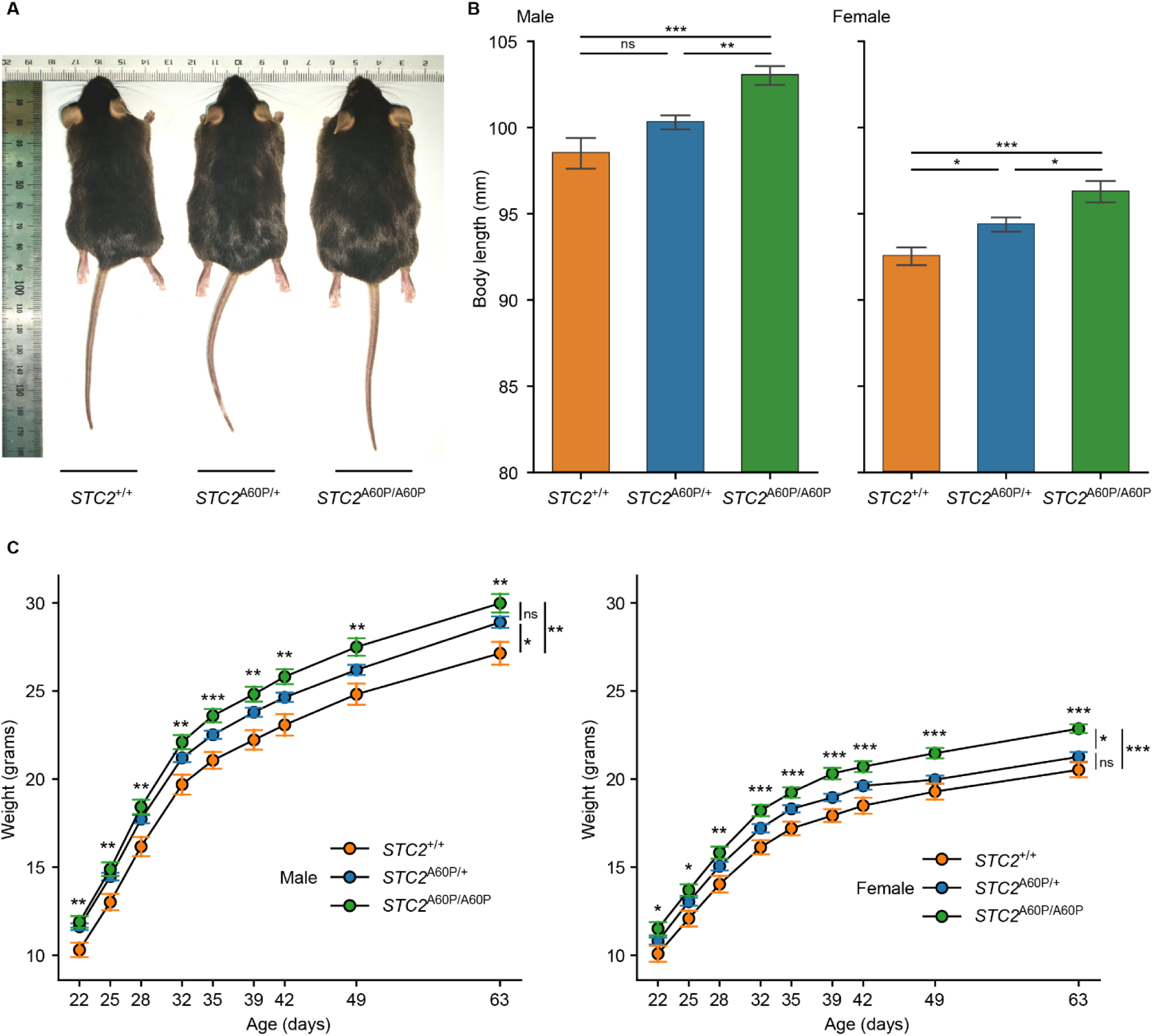
STC2 A60P increase body weight and body length in mice. **(A)** Representative image of littermate STC2 A60P gene edited mice. **(B)** Body length of *STC2*^+/+^, *STC2*^A60P/+^, *STC2*^A60P/A60P^ mice at 14 weeks of age. Male n = 12, 40, 17; Female n = 12, 30, 13. Error bars and asterisks as in Fig. 3B. One-way ANOVA followed by Tukey’s post hoc multiple comparison test was used. **(C)** Total body mass of indicated littermates from 3 to 9 weeks of age. Male n = 12, 40, 17; Female n = 12, 30, 13. Error bars and asterisks as in Fig. 3A. Two-sided Student’s t-tests were used to infer significant body mass differences between *STC2*^A60P/A60P^ and *STC2*^+/+^ mice at each time point (notations above each time point). Subsequently, repeated measures two-way ANOVA followed by Tukey’s post hoc multiple comparison test was conducted to infer significant body mass differences among the three genotypes (notations on the far right).

Consistent with our expectations, the STC2 A60P mutation also demonstrated an additive effect on body size increase (Fig. 6B). At 14 weeks of age, *STC2*^A60P/+^ males and females were respectively 1.8% (1.8 mm) and 2.0% (1.8 mm) longer than *STC2*^+/+^ mice, whereas *STC2*^A60P/A60P^ males and females were respectively 4.6% (4.5 mm) and 4.0% (3.7 mm) longer than *STC2*^+/+^mice.

### Distribution and evolution of rs384548488*A, rs110540352*G, and rs42661323*G in cattle

We investigated the origin and distribution of rs384548488*A, rs110540352*G, and rs42661323*G mutations in ancient and modern cattle. The frequencies of these three mutations in different breeds were calculated by using run 9 of the 1000 Bull Genomes (Table S12). Most of the 478 *Bos indicus* individuals lack these three mutations except for Brahman cattle, a cross of *Bos indicus* and *Bos taurus*. Among 225 Brahman cattle, only one rs384548488*A, two rs110540352*G, and three rs42661323*G alleles were found. This suggests all three mutations emerged after *Bos taurus* and *Bos indicus* divergences.

For the *LCORL* locus, breeds with rs384548488*A frequencies > 0.5 predominantly originated in the Alpine region (Fig. 7A). The rs384548488*A allele was nearly fixed in Original Braunvieh (frequency=0.992) and the related Brown Swiss (frequency=0.997) and in several other central European breeds. Remarkably, the allele is almost absent in black Angus (frequency=0.014) but has a clearly higher frequency in red Angus (frequency=0.386). However, the rs384548488*A allele was not detected in 16 ancient Middle Eastern cattle spanning 655- 8123 years ago (Fig. 7B). This suggests that rs384548488*A likely emerged after the spread of domestic *Bos taurus* from the Fertile Crescent to Europe, possibly originating in cattle of Braunvieh ancestry in the Alpine region.

**Fig. 7.**
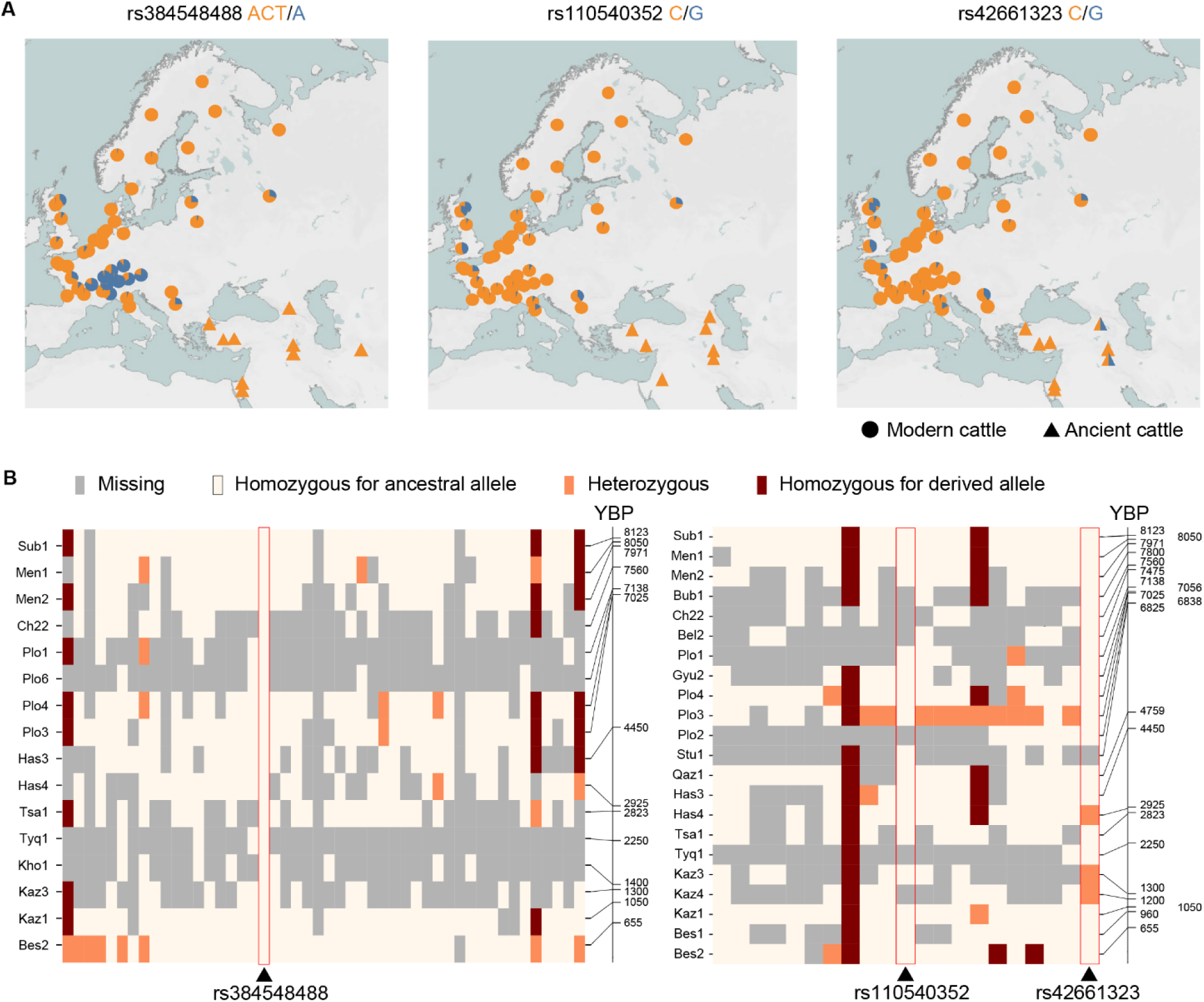
Distribution and Evolution of *LCORL* and *STC2* variant in Cattle. **(A)** Geographical distribution of *LCORL* variant rs384548488*A, and of *STC2* variant rs110540352*G and rs42661323*G. Blue and orange indicate the derived alleles and ancestral alleles, respectively. The triangles half orange and half blue indicate that this ancient cattle is a heterozygote. The modern cattle data is from in run 9 of the 1000 Bull Genomes. **(B)** Genotypes of *LCORL* and *STC2* locus in ancient cattle. The triangles indicate the casual variants. The missing genotypes are colored in grey, homozygotes for ancestral allele are light-yellow, heterozygotes areorange, homozygotes for derived allele are dark-red.

At the *STC2* locus, the frequencies of both causal variants, rs110540352*G and rs42661323*G, were below 0.5 in all breeds (Fig. 7A). Hereford has the highest frequencies of rs110540352*G and rs42661323*G. In Alpine cattle breeds where the rs384548488*A frequency exceeded 0.5, the maximum frequencies of rs110540352*G and rs42661323*G were only 0.077, whereas the derived alleles are absent in Original Braunvieh and Brown Swiss. According to ARG, the inferred age of rs110540352*G in five beef cattle populations at 911 years to 1,148 years, which is later than the inferred age of rs42661323*G at 6,571 years to 9,761 years.

Consistent with this, rs110540352*G was not present in ancient cattle samples, while rs42661323*G was identified as heterozygous in one Iron Age and two Middle Ages ancient cattle samples (Fig. 7B). Although rs110540352*G emerged later than rs42661323*G, its frequency in modern cattle is slightly higher than that of rs42661323*G. Selection coefficients in Hereford inferred by CLUES also indicate that rs110540352*G (s=0.012) has been under stronger selection pressure than rs42661323*G (s=0.007) in the last 1000 years.

## DISCUSSION

Previous GWASs have identified a set of genes as shared regulators of body size in domesticated animals ^2,3^. However, the high LD patterns posed challenges for pinpointing causal variants. The ARG encompasses the complete history of genomic mutations, facilitating detailed and robust inferences about selection ^36,37^. The CLUES method, based on the ARG, has shown excellent power in inferring recent selection and fine-mapping selected sites ^19^. In this study, we utilized these methods to identify 11 loci driving recent selective sweeps in body size QTLs between British and Continental beef cattle. Specifically, we proposed sweep-driving mutations at the top effect loci, *LCORL* and *STC2*. Then, we validated causal coding mutations on *LCORL* and *STC2* in gene-editing mice, which support the role of these mutations in growth. This marks the first time that causal variants regulating body size and growth traits in cattle have been experimentally verified *in vivo*. We propose that the utilization of ARG and the CLUES method is an effective tool for fine-mapping recently selected sites in economic traits QTLs of domesticated animals.

The *LCORL* locus is reported as being associated with body size in various domesticated animals, such as dogs ^3^, horses ^31^, goats ^30^, and cattle ^2,8–15,38^. In this study, we propose that pLoPD mutations are putative sweep-driving alleles in *LCORL* in sheep, pigs, rabbits, and chickens. These results highlight its role as a convergent sweep-driving mutation across above species. Mouse models confirmed the growth-promoting effects of the pLoPD mutation in PALI2. It is important to note that PALI2 is only one type of *LCORL* expression product, and complete *LCORL* knockout in mice leads to different outcomes ^39^. The possible function of PALI2 is to enhance PRC2-mediated methyltransferase activity ^22^. In mouse embryonic stem cells, PRC2 targets a range of well-known developmentally regulated genes, including the *Wnt6-Ihh* and *Hoxb* loci ^25,40^. We found that the loss of the PIP domain in PALI2 has a more pronounced effect on H3K27me3 levels during embryonic development than in adult mice. This underscores the significance of PRC2-mediated methyltransferase activity in growth regulation, suggesting a particularly strong role during embryonic development.

The IGF system plays a central role in regulating overall body growth by controlling cellular growth, differentiation, and survival ^41^. Consequently, numerous genes within the IGF pathway, including *IGFs*, *IGFBPs*, *GHR*, and *STC2*, are crucial for growth regulation in both humans and domesticated animals ^2,3,21,42,43^.

A study on human height showed that two rare variants in human STC2, p.R44L and p.M86I, disrupt the inhibition of PAPP-A-mediated proteolysis of IGFBP-4 in vitro, significantly increasing height by 1.9 cm and 0.9 cm, respectively ^33^. In cattle, we identified a causal missense mutation, rs42661323G (STC2 p.A60P), located within the interaction region between STC2 and its ligand PAPP-A. Consequently, saturation mutagenesis on the STC2-PAPP-A interaction domain via gene editing emerges as a promising strategy for generating novel potentially, advantageous STC2 alleles. Additionally, we identified a selective noncoding mutation, rs110540352G, downstream of the *STC2*, which is located in active regulatory elements in the cerebral cortex and hypothalamus. Due to the high linkage disequilibrium between rs110540352*G and rs42661323*G, haplotypes carrying both variants are expected to have a combined effect of reducing *STC2* expression and producing functionally impaired STC2 protein.

The causal mutations in *LCORL* and *STC2* identified in this study are naturally occurring and are present high allele frequency in Britain or central-Europe cattle. The pLoPD mutation in *LCORL* is also common in multiple domestic animals, and in wild rabbits and wild chickens. The amino acid substitution p.A60P in STC2, which is highly conserved among vertebrates, also may be a potentially useful target for genetic improvement. In summary, the mutations identified in this study that regulate body size and weight traits in cattle provide precise targets for genome selection and gene editing, potentially facilitating the enhancement of economically important traits in all kinds of livestock species.

## RESOURCE AVAILABILITY

### Lead contact

Requests for further information and resources should be directed to and will be fulfilled by the lead contact, Yu Jiang.

### Materials availability

This study did not generate new unique reagents.

### Data and code availability

- Relate-estimated coalescence rates and allele ages for the Charolais, Simmental, Limousin, Hereford, Angus cattle have been deposited at Zenodo and are publicly available at https://zenodo.org/records/14259711. Our predictions of cattle ancestral alleles for SNVs have been deposited at Zenodo and are publicly available at https://zenodo.org/records/14261687.
- All original code has been deposited at GitHub repository (https://github.com/Bai-fengting/cattleARG_LCORL_STC2)
- Any additional information required to reanalyze the data reported in this paper is available from the lead contact upon request.

## Materials and Methods

### Variant processing from the 1000 Bull Genomes Project Run 9 for ARG construction

We utilized bcftools version 1.16 ^44^ for variant filtering. We excluded variants falling within the 99.90 to 100.00 tranche of the GATK Variant Quality Score Recalibration (VQSR) for SNPs. Only biallelic SNPs with a missing rate of less than 20% were retained. To infer the ARG, we first removed singleton variants and then applied Beagle version 5.4^45^(with parameters: window=5, overlap=2) for phasing the remaining variants.

### Individual selection for ARG construction

Based on the sample information provided by Run 9 of the 1000 Bull Genomes Project, the sample sizes for Angus, Hereford, Charolais, Simmental, and Limousin were 401, 142, 154, 137, and 101, respectively. To construct the ARG, we aimed to select 100 individuals from each breed.

We used KING 2.3.0 ^46^ to calculate kinship coefficients between individuals (–kinship). To retain a larger number of unrelated individuals within each breed, we employed the following strategies:

For Angus, Hereford, and Charolais:

We iteratively removed individuals with the most third-degree or closer relationships (kinship coefficient > 0.0442) within each breed until all remaining individuals were unrelated. We then retained the 100 individuals with the highest sequencing depth.

For Simmental:

We iteratively removed individuals with the most third-degree or closer relationships (kinship coefficient > 0.0442) within each breed until all remaining individuals were unrelated. This process yielded 81 Simmental individuals. We applied the same method to select 19 German Simmental (Fleckvieh) individuals, which were then combined with the Simmental group to reach a total of 100.

For Limousin:

Due to the limited sample size of 101, we only removed the individual with the lowest sequencing depth, retaining 100 individuals.

In addition to these 500 individuals from the five main breeds, we selected 168 individuals from other breeds to jointly infer the preliminary ancestral recombination graph.

### Determination of bovine ancestral alleles

We retained 2,144 individuals with BioProject IDs from Run 9 of the 1000 Bull Genomes Project. To obtain a representative sample, we selected one individual with the highest sequencing depth from each breed, resulting in a total of 79 representative individuals.

The assignment of bovine ancestral alleles was based on a model comparison of alleles from cattle with alleles from outgroup species: Water Buffalo, Sheep, and White-Tailed Deer. While species of the Bos genus are more closely related and potentially more informative, they share a high proportion of genetic introgression with cattle ^47^.

We utilized multiple sequence alignments of 110 species (78 ruminants and 32 mammalian outgroup species), available from http://animal.omics.pro/code/index.php/RGD/loadByGet?address[]=RGD/Download/comSynDo <u>wnload.php</u> ^48^, to determine the alleles in Water Buffalo, Sheep, and White-Tailed Deer at each locus. We employed MafFilter v1.3.1^49^ to generate sequence alignment files between cattle and each of the three outgroup species. We then used our custom script to retrieve sequence data for cattle and the three outgroup species in est-sfs input format. Only sites with sequence data available in at least one outgroup species were retained, resulting in 72,831,650 sites across the four species used for ancestral allele determination.

We employed the est-sfs software ^50^ with the K2 model to infer the probability (Pancs) of the major allele in cattle being ancestral. Alleles were determined to be ancestral if they were the major allele at a site with Pancs > 0.8 or the minor allele at a site with Pancs < 0.2. For the remaining sites where ancestral alleles could not be determined, we used the reference allele as the ancestral allele, as Relate software ^18^ has the capability to adjust ancestral alleles during analysis.

### Constructing ARG and estimating relative coalescence rates through time

We constructed the ARG using Relate version v1.1.9 ^18^, applied to 689 phased whole-genome sequences. The following parameters were used: mutation rate: 1.26 × 10 per base per generation; recombination rate: 1.00 × 10; effective population size (Ne): 10,000; seed: 1. Subsequently, we used Relate’s EstimatePopulationSize.sh script to estimate coalescence rates and re-estimate branch lengths. The parameters for this step were: mutation rate: 1.26 × 10; generation time: 6 years; number of iterations: 10; tree dropping threshold: 0.5; painting: 0.025 1; time bins: defined as 10 years ago, where x ranges from 2 to 6.6 with an increment of 0.1.

To infer relative cross-coalescence rates between the five beef cattle breeds, we extracted subtrees such that all tips belonged to these five breeds and re-estimated coalescence rates. We then extracted coalescence rates and cross-coalescence rates from the.coal files to calculate relative cross-coalescence rates.

### Fst and CLUES analyses

We grouped Angus and Hereford as one population and Charolais, Simmental, and Limousin as another population. Using vcftools 0.1.16 ^51^, we performed Fst analysis on each SNV in the ARG for these five beef cattle breeds. The top 0.05% of SNVs with the highest genome-wide Fst values were considered highly differentiated and selected for subsequent CLUES analysis. To reduce false positives caused by potentially inaccurate ARG inference in some local regions, we applied the following filters to remove SNVs: 1. SNVs with the number of mutations mapping to their tree in the bottom 5th percentile; 2. SNVs where the fraction of tree branches having at least one SNP is in the bottom 5th percentile.

We extracted ARG of each breed separately and re-estimated coalescence rates and branch lengths as input for the CLUES selection test ^19^. For each highly differentiated SNV, we first used Relate’s SampleBranchLengths.sh to extract the corresponding local tree and resampled branch lengths 200 times. CLUES calculates a likelihood ratio for each input SNV, reflecting the degree to which the derived allele of this SNV deviates from the neutral mutation model. We repeated the calculation three times for each highly differentiated SNV using different seeds (seed = 1, 10, 100). The median of the log(LR) values was then assigned to each SNV.

### CLUES selection test adjusted for genetic drift in population history

To reduce false positives caused by genetic drift and determine the threshold for rejecting neutral mutations in CLUES analysis, we simulated a series of neutral mutations based on the effective population size changes over time, as estimated by Relate for the five beef cattle breeds. We calculated the changes in effective population size over time based on the coalescence rates estimated by Relate for the five beef cattle breeds. As all five beef cattle breeds are of European origin, their effective population size trends were generally consistent. We used the average effective population size of the five breeds for subsequent data simulation. The msprime 1.2.0 ^52^was applied to simulation with parameters: sequence length: 158,534,110 bp (length of chromosome 1 in ARS-UCD1.2); number of individuals: 100 (200 haplotypes), matching each beef cattle breed; recombination rate: 1 × 10; mutation rate: 1.26 × 10; random seed: 1; model: Classical coalescent with recombination model (Hudson’s algorithm) We converted all simulated biallelic SNVs into Relate input format and inferred the ancestral recombination graph using the same parameters as for the cattle ARG construction. The effective population size changes inferred from the simulated data showed the same trend as the real data within ∼300,000 years (the divergence time between taurine and zebu cattle). After removing low-quality sites, we obtained 1,006,489 SNVs. We selected one SNV every 100 positions, resulting in 10,064 SNVs for the CLUES selection test, following the same procedure as for the real data. The resulting log(LR) distribution from simulated data, with 95%, 99%, and 99.9% percentiles, is shown in Fig. S2.

### Genome coordinate remapping

To ensure consistency with our analysis based on the ARS-UCD1.2 reference genome, we performed coordinate remapping for two sets of previously published data. First, we addressed the GWAS result for daily weight gain in beef cattle conducted by Zhang et al ^12^. In their study, the physical positions of variants were based on the UMD3.1 reference genome. To align these coordinates with our current framework, we utilized NCBI’s Remap Tool to reposition the UMD3.1-based physical coordinates onto the ARS-UCD1.2 reference genome.

Secondly, we focused on the cattle stature QTLs reported by A. C. Bouwman et al ^2^. Similar to the previous dataset, the genomic coordinates for these QTLs were originally based on the UMD3.1 reference genome. For this remapping, we employed the UCSC LiftOver tool ^53^ to translate the UMD3.1-based physical coordinates to their corresponding positions on the ARS- UCD1.2 reference genome.

### Mapping and variant calling for ancient cattle genome data

Our ancient cattle genome data were sourced from the study by Verdugo et al ^54^. We followed their methodology for processing the ancient cattle genome sequencing data, with some modifications. The specific steps were as follows:

We first used cutadapt ^55^ to remove adapters from reads and filter out low-quality reads, using the following parameters:-a AGATCGGAAGAGCACACGTCTGAACTCCAGTCAC-O 1-m 30. We then aligned the quality-controlled reads to the ARS-UCD1.2 reference genome using the aln module of bwa 0.7.17^56^, and used the samse module with the option-r for defining read groups and producing unfiltered SAM files. We used samtools 1.14 ^57^ for quality control and sorting of reads in the SAM files, generating BAM files (-Sb-F 4-q 25), followed by duplicate removal. Picard tools 2.16.0 was used to merge BAM files. Indel realignment was performed using GATK 3.8-0-ge9d806836 ^58^. For samples Th7 and Ch22, we removed duplicates using Picard tools 2.16.0 before performing indel realignment.

For all autosomes, we used bcftools’s mpileup and call modules (bcftools call-m) to call SNVs on the filtered ancient cattle BAM files, based on the positions of biallelic SNVs from the 1000 Bull Genomes project. The variant rs384548488 (Chr6:g.37401771_37401772del) is an INDEL. To obtain the genotypes of rs384548488 and nearby SNVs in ancient cattle, we performed genotyping on all sites in the genomic region chr6:37151770-37651770 using bcftools’s mpileup and call modules (bcftools call-m).

### Mapping and variant calling for modern genome data

For the downloaded FASTQ files, we used fastp 0.23.4 ^59^ with default parameters to remove adapters and low-quality reads. The processed reads were then analyzed following the GATK Best practices for data pre-processing for variant discovery and somatic short variant discovery (SNVs + Indels). In brief, we first used the MEM module of BWA to map reads to the reference genome. Subsequently, we used samtools to sort the BAM files and picard tools to remove duplicates. GATK HaplotypeCaller, CombineGVCFs, and GenotypeGVCFs were employed for joint genotyping of multiple samples.

### Prediction of variant effect

To annotate variants in cattle, we utilized VEP (Variant Effect Predictor ^60^ from Ensembl release 112 and cache version 110 to predict variants effect. We assigned 5 kb as the distance up and/or downstream between a variant and a transcript for which VEP will assign the upstream_gene_variant or downstream_gene_variant consequences. Additionally, we employed ANNOVAR ^61^ to annotate variants located in intergenic regions. We defined intergenic regions as being the same if they share the nearest upstream and downstream genes.

### Scanning pLoPD variants

To identify potential variants causing the loss of the PIP domain in PALI2, we first obtained the amino acid sequences of the PIP domain for cattle, rabbit, and chicken PALI2 from the study by E. Conway et al ^22^. For sheep, goat, horse, dog and pig, where known PALI2 PIP domain sequences were unavailable, we used the cattle PALI2 PIP domain sequence as a proxy. We then employed NCBI’s online BLAST tool to align these PIP domain amino acid sequences to the reference genomes of sheep and pig, thereby determining the genomic coordinates of the PALI2 PIP domain in these species. The reference genomes used for cattle, sheep, goat, pig, horse, dog, rabbit, and chicken were ARS-UCD1.2, ARS-UI_Ramb_v2.0, ARS1, Sscrofa11.1, Equcab3.0, ROS_Cfam_1.0, OryCun2.0, and GRCg7w, respectively. For gene annotations, we utilized Ensembl release 104 for ARS-UCD1.2 and ROS_Cfam_1.0 and Ensembl release 112 for ARS- UI_Ramb_v2.0, ARS1, Sscrofa11.1, and OryCun2.0. As Ensembl annotations were not available for the GRCg7w genome, we used NCBI gene annotations for chicken.

Rice et al. published a pangenome reference (GRCg7b) constructed from 30 chickens^28^.

However, neither Ensembl nor NCBI gene annotations for GRCg7b fully annotated the mRNA and exons of *LCORL* containing the PIP domain. Consequently, we couldn’t directly assess the impact of variants on the PALI2 PIP domain based on GRCg7b annotations. Therefore, we aligned the genomic coordinates of variants based on GRCg7b to GRCg7w using BLAST, allowing us to evaluate the potential effects of these variants on the PALI2 PIP domain based on NCBI gene annotations for GRCg7w genome.

For sheep, pig, and rabbit, we obtained the ancestral sequences near the *LCORL* frameshift intron from the Ensembl 43 eutherian mammals EPO multiple alignment data ^62^.

### Haplotype and genotype pattern analysis

For cattle, to analyze the haplotype patterns composed of both INDELs and SNVs, we employed bcftools version 1.16 to filter variants. We retained SNPs and INDELs with a missing rate of less than 20%. Variants from the GATK VQSR (Variant Quality Score Recalibration) 99.90 to 100.00 Tranche for both SNPs and INDELs were excluded from our analysis. We then used beagle5.4 with parameters window=10 and overlap=2 to phase the SNPs and INDELs.

For pig, sheep, we applied the GATK-recommended hard filtering thresholds to filter INDELs and SNVs. Specifically, for SNVs, we used the following criteria: INFO/QD >= 2.0 & QUAL >= 30.0 & INFO/SOR <= 3.0 & INFO/FS <= 60.0 & INFO/MQ >= 40.0 & MQRankSum >=-12.5 & INFO/ReadPosRankSum >=-8.0. For INDELs, we applied these thresholds: INFO/QD >= 2.0 & QUAL >= 30.0 & INFO/FS <= 200.0 & INFO/ReadPosRankSum >=-20.0.

After filtering, we used beagle5.4 (window=10, overlap=2) to phase SNPs and INDELs for pig and sheep variants. For all species studied, we extracted mutations within the *LCORL*. To visualize the haplotype or genotype patterns, we developed custom scripts to generate heatmaps. The bifurcation diagram for haplotypes is plotted using R package rehh^63^.

### Motif prediction

To predict transcription factor binding sites, we employed the MAST (Motif Alignment and Search Tool) from the MEME Suite 5.5.5 ^64^. Our approach began with the extraction of the sequence of the active regulatory element (Chr20:4967400-4969800) containing rs110540352 from the reference genome using samtools 1.16. We then utilized the PFM (Position Frequency Matrix) data for vertebrates from the JASPAR CORE database 2024 ^65^ as our motif database.

Using MAST with its default parameters, we scanned the extracted sequence for potential transcription factor binding sites. This analysis was performed twice: once for the sequence containing the ancestral allele at rs110540352, and once for the sequence containing the derived allele. By comparing the MAST results for both the reference and alternate sequences, we were able to predict how the rs110540352 variant might affect transcription factor binding within this active regulatory element.

### Luciferase assays

The ancestral *STC2* regulatory element fragment (AAGTGGCGCCTCAA), the mutational *STC2* regulatory element fragment (AAGTGGCGCCTGAA), and the E2F4 coding sequence(ENSBTAT00000015999.7) synthesized by Sangon Biotech (Shanghai, China). The E2F4 coding sequence was inserted into pCDNA3.1_3XFlag (Addgene, #182494) named pCDNA3.1_3XFlag-E2F4. The ancestral *STC2* regulatory element fragment and the mutational *STC2* regulatory element fragment were inserted into the luciferase reporter vector pGL4.10 named pGL4.10-ANC and pGL4.10-MUT, respectively.

HEK293T cells (ATCC) were cultured in DMEM (Gibco) supplemented with 10% FBS (Gibco) and incubated at 38.5°C in a 5% CO2 environment. Approximately 1 μg of plasmid DNA (0.45 μg each for pGL4.10-ANC or pGL4.10-MUT, 0.45 μg for pCDNA3.1_3XFlag-E2F4, and 0.1 μg for pRL-TK) was cotransfected using PEI Transfection Reagent (MCE) according to the manufacturer’s protocol. The pRL-TK plasmid served as an internal control for normalizing transfection efficiency. Cell lysates were collected 48 hours post-transfection and prepared for luciferase activity analysis using the Double-Luciferase Reporter Assay Kit (TransGen Biotec) following the manufacturer’s instructions. Relative luciferase activities were expressed as the ratio of firefly luciferase to Renilla luciferase values.

### Quantification of allele expression

In Hereford cattle, rs110540352*G and rs42661323*G are in high linkage disequilibrium. The rs110540352(C>G) is located in an intergenic region, while rs42661323(C>G) is situated in an exon of the *STC2*. To investigate the effect of rs110540352(C>G) on *STC2* expression in the hypothalamus and cerebral cortex, we counted the reads carrying rs42661323*C and rs42661323*G in individuals heterozygous for rs42661323. By analyzing the potential differences between reads carrying rs42661323*C and rs42661323*G, we aimed to infer the impact of rs110540352(C>G) on *STC2* expression.

We selected RNA-seq data from the cerebral cortex of Hereford cattle from the CattleGTEx dataset. To process this data, we used fastp 0.23.4 with parameters of cut_window_size 4 and cut_mean_quality 15 to remove adapters and low-quality reads. Subsequently, we employed Hisat2 ^66^ with default parameters to map the RNA-seq reads to the ARS-UCD1.2 reference genome and used samtools 1.16 to sort the reads. We further utilized Picard toolkit 3.0.0 to remove duplicates.

For visualization and quantification, we used IGV 11.0.1 ^67^ to inspect the bam files and count the reads carrying rs42661323*C and rs42661323*G. Due to the relatively low number of reads obtained for each sample, we combined the counts of reads carrying rs42661323*C and rs42661323*G from all samples for our analysis.

### Predicting the effect of STC2 A60P

To predict the effect of the STC2 A60P mutation, we employed a multi-faceted approach combining comparative genomics, structural biology, and computational prediction tools. We obtained multi-sequence alignment results from the UCSC Multiz Alignments of 100 Vertebrates for comparative genomic analysis. This alignment provided a broad evolutionary perspective on the conservation of the STC2 A60 across vertebrate species.

For structural analysis, we utilized the human PAPP-A·STC2 complex structure published by Kobberø et al^35^. We downloaded this structural data and used PyMOL 2.5.8 ^68^ for three-dimensional protein structure visualization. This allowed us to examine the spatial relationship between STC2 and PAPP-A, and to visualize the potential impact of the A60P mutation on this interaction.

To assess the physicochemical properties of the mutation, we employed ProtScale (https://web.expasy.org/protscale/) to predict the hydrophobicity of STC2 before and after the A60P mutation. This analysis provided insights into how the mutation might alter the local environment of the protein and potentially affect its function or interactions.

Furthermore, we used BeAtMuSiC v1.0 ^69^ to predict the impact of STC2 A60P on the binding affinity between STC2 and PAPP-A. This computational tool allowed us to estimate the change in binding free energy caused by the mutation, providing a quantitative prediction of how the A60P substitution might affect the protein-protein interaction.

### Validating the function of LCORL lacking the PIP domain in mice

We performed sequence alignment of the exons containing the PIP domain of *LCORL* in cattle and mice using the MUSCLE algorithm in MEGA 11 ^70^. This analysis identified the homologous position of the cattle rs384548488 variant in mice as Chr5:45,882,519-45,882,520 (Fig. S15A). We targeted the region Chr5:45,882,459-45,882,558 for CRISPR/Cas9-mediated deletion to induce a frameshift mutation, which was carried out by Cyagen Biosciences Inc (Suzhou, China). Animal experiments were performed in accordance with the regulations and guidelines established by the Animal Care Committee of Northwest A&F University. Briefly, gRNAs targeting the mouse *LCORL* (gRNA-A1 matching the reverse strand: AGGGTTAAAAGATTCATTTTGGG, and gRNA-A2 matching the forward strand: TTGTCACTGTTGTTTATGGAAGG) were co-injected with Cas9 mRNA into fertilized C57BL/6J mouse oocytes to generate offspring with the targeted gene deletion. F0 founder animals were identified through PCR (Primers: F1: 5’-CAAGAAGACCCTAAGGAAAAGTCA-3’, R1: 5’-TCTGAGGTATCATAGACTTGCTCT-3’) followed by sequencing analysis. These animals were subsequently mated with wild-type mice to test germline transmission and produce F1 generation animals.

We commissioned Cyagen Biosciences Inc (Suzhou, China) to perform in vitro fertilization (IVF) techniques using heterozygous male and female mice to rapidly generate the F2 generation (N=112; 56 males, 55 females). All mice were maintained under ad libitum feeding conditions, with no more than 5 mice per cage. We commissioned Cyagen Biosciences Inc (Suzhou, China) to measure the body weight of 10 individuals per genotype for both male and female mice, starting from weaning (3 weeks of age). All mice undergoing weight measurements were housed in groups of 5 per cage. All mice were kept under identical environmental conditions with free access to food. Inferences about differences between means were tested by two-tailed Student’s t-test.

### Validating the function of STC2 A60P in mice

We commissioned Cyagen Biosciences Inc (Suzhou, China) to generate STC2 A60P gene-edited mice. Animal experiments were performed in accordance with the regulations and guidelines established by the Animal Care Committee of Northwest A&F University. Briefly, a gRNA targeting the mouse Stc2 gene (matching forward strand of gene: CAGCACTGTTTGGTCAATGCCGG), a donor oligonucleotide containing the p.A60P (GCC to CCC) mutation, and Cas9 were co-injected into fertilized C57BL/6J mouse oocytes to produce offspring with the targeted gene knock-in. F0 founder animals were identified through PCR (Forward primer (F1): 5’-CGATAGAAGGAAGAAAAGAAAACGC-3’, Reverse primer (R1): 5’-CAAACACCAGACTCTCTCAAGCAA-3’) followed by sequence analysis. These animals were subsequently mated with wild-type mice to test germline transmission and produce F1 generation animals. We commissioned Cyagen Biosciences Inc (Suzhou, China) to use in vitro fertilization (IVF) techniques with heterozygous male and wild-type female mice to generate 30 heterozygous male and 30 heterozygous female mice rapidly. We then bred these 30 pairs of heterozygous mice to produce the F2 generation (N=126; 70 males, 56 females). Body weight measurements were conducted on 125 mice. All mice were maintained under identical environmental conditions with free access to food, housed in groups of 5 per cage. Inferences about differences between means were tested by two-tailed Student’s t-test.

### Total protein extraction and western blotting

First, the embryos, thymuses and muscles of control and mutant groups were washed with ice-cold PBS. Then added 3mm stainless steel beads and the RIPA Lysis Buffer (Solarbio, Beijing, China) mixed with 1% protease inhibitor (05056489001, Roche, Switzerland) and 1mM PMSF (ST506, Beyotime, Shanghai, China) to grind for extracting total proteins.

After measuring protein concentration with BCA kit (ZJ102, Epizyme, Shanghai, China), the protein samples were boiled in 1×Protein Loading Buffer (LT101S, Epizyme, Shanghai, China) to denaturate all the proteins and separated by 10% SDS-PAGE gel (PG212, Epizyme, Shanghai, China). Proteins were transferred to PVDF membranes by wet transfer, and blocked with 5% nonfat milk for 2h. Then, the membranes were incubated with primary antibody at 4 overnight, followed by incubation with HRP-coupled secondary antibody at room temperature for 2h.

Finally, blots were visualized with a chemiluminescence detection system. The antibodies used are as follows: H3K27me3 (9733s, Cell Signaling Technology, 1:1000), β-actin (8457, Cell Signaling Technology, 1:1000), β-tubulin (2128S, Cell Signaling Technology, 1:1000).

## ACKNOWLEDGMENTS

We thank Wen Wang, from School of Ecology and Environment, Northwestern Polytechnical University, China, for suggestions and comments.

We thank the High-Performance Computing platform of Northwest A&F University and Computing Center in Xi’an for providing computing resources.

The project was supported by National Key R&D Program of China (2022YFF1000100, 2023YFD1300402), National Natural Science Foundation of China (U21A20120), Shaanxi Livestock and Poultry Breeding Double-chain Fusion Key Project (2022GD-TSLD-46-0401).

## AUTHOR CONTRIBUTIONS

Conceptualization: Y.J., Y.C., F.B.

Data curation: F.B., S.J.

Formal analysis: F.B., Y.C., M.Q.

Funding acquisition: Y.J., X.W.

Investigation: L.P, Y.L, C.L., Y.F., S.G., M.L.

Methodology: Y.C., F.B., Y.J.

Project administration: Y.J.

Resources: Y.J., G.R.

Software: F.B., Y.C.

Supervision: Y.J.

Validation: Y.J., G.R.

Visualization: F.B., M.Q., Q.Y.

Writing – original draft: F.B.

Writing – review & editing: Y.J., Y.C., R.H., J.A.L.

## DECLARATION OF GENERATIVE AI AND AI-ASSISTED TECHNOLOGIES

During the preparation of this work the authors used ChatGPT in order to improve the readability and language of the manuscript. After using this tool, the authors reviewed and edited the content as needed and take full responsibility for the content of the published article.

