## Supplemental Figure 1-15 Supplement Table 6-9, 11 for "*LCORL* and *STC2* variants increase body size and growth rate in cattle and other animals"

Fengting Bai *et al.*

**This PDF file includes:**

Figs. S1 to S14

Tables S6 to S9, S11

Legends for tables S1 to S5, S10, and S12

**Other Supplementary Materials for this manuscript include the following:**

Tables S1 to S5, S10, and S12

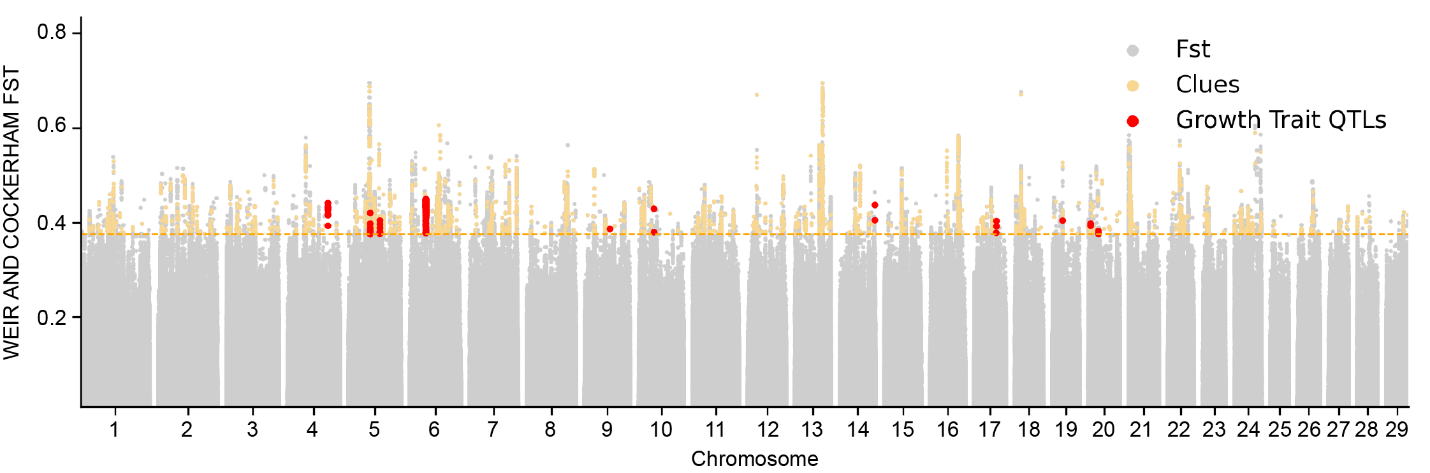

Fig. S1.

**Genome-wide Fst analysis between British and Continental European breeds and CLUES selection test in the past 1,000 years.** Scatter points represent *Fst* values of genome-wide SNVs between British and Continental European breeds. The dashed line indicates the genome-wide top 0.05% threshold. Yellow dots represent loci that have been under positive selection in at least one breed in the past 1,000 years. Red dots indicate loci located within 11 cattle body size QTLs.

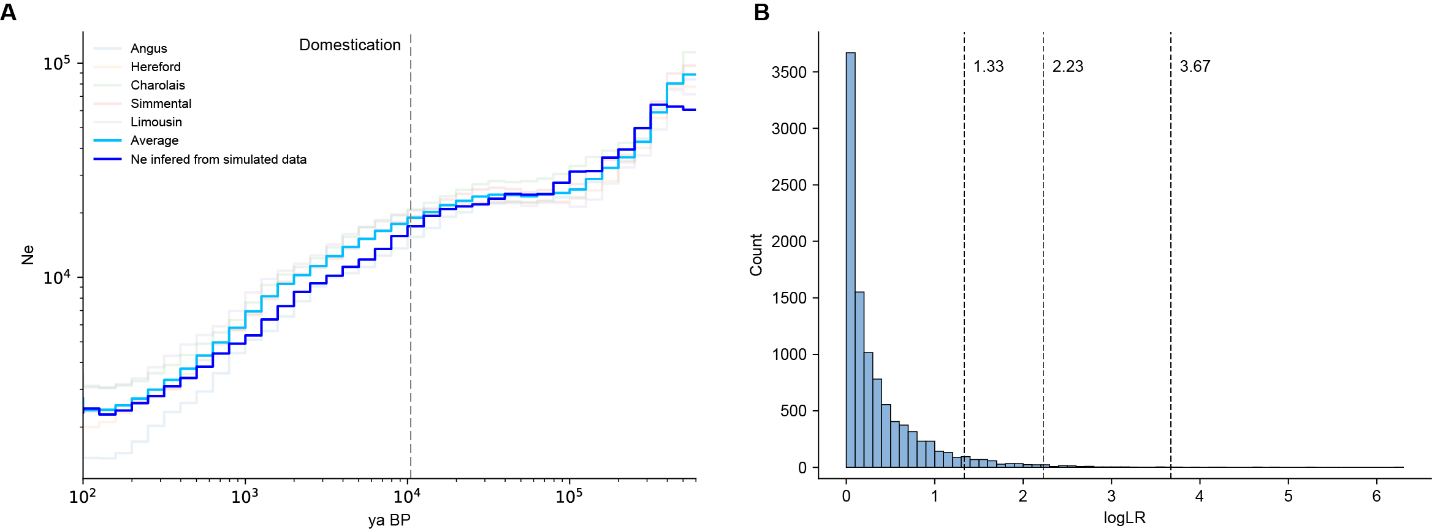

Fig. S2.

**Establishing the logLR threshold for rejecting neutrality.** **(A)** Population size history inference of five European beef cattle breeds using Relate. The line in light blue shows the average population size history. The line in light blue shows population size history inferred from simulated data using Relate. **(B)** Distribution of logLR from 10064 simulated datasets. The three vertical lines in light blue show 95%, 99%, and 99.9% percentile points, respectively. We use the 95% percentile point of 2.23 as our neutrality rejection threshold.

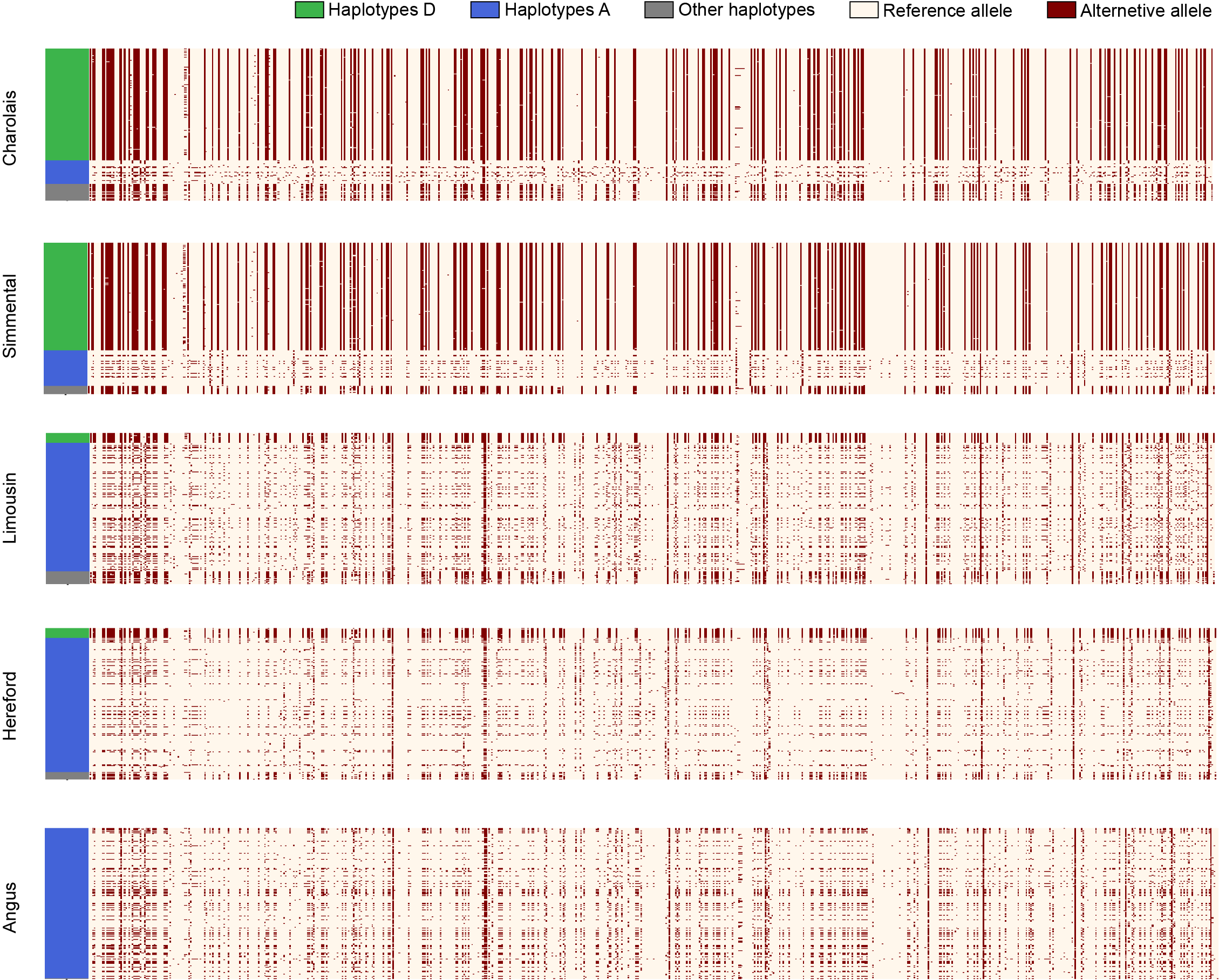

**Fig. S3.**

**Haplotype Patterns at the selection-targeted *NCAPG-LCORL* region (Chr6: 37349373-37438248).** Haplotypes are ordered as follows: haplotypes D, haplotypes A, and other haplotypes. The D haplotypes are mainly found in Charolais and Simmental cattle, but not in the Angus. Each row represents a haplotype, and each column represents a SNV. The positions along the x-axis are shared between panels A and B.

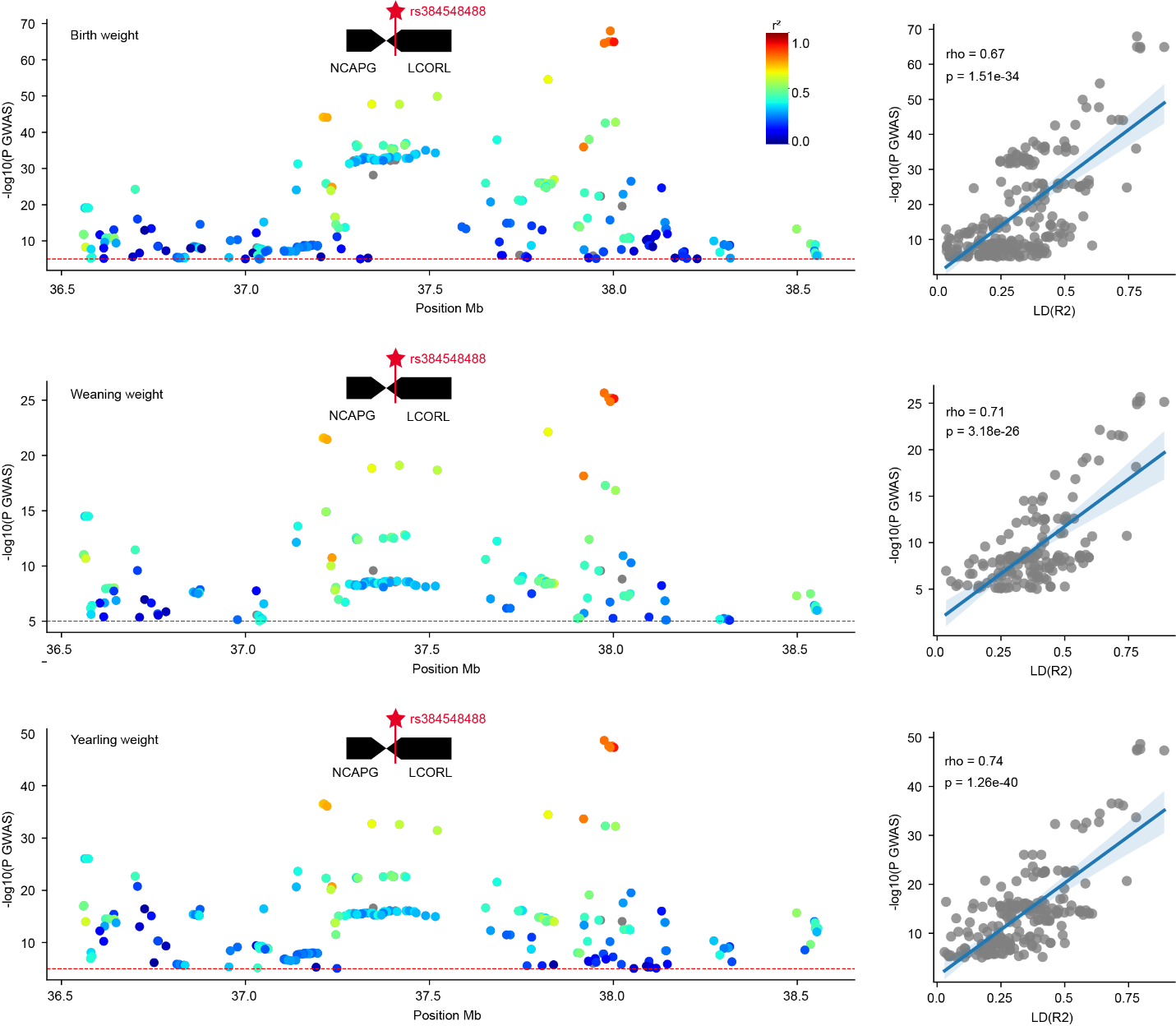

Fig. S4.

**The rs384548488 variant is associated with growth trait variation in Red Angus cattle.** The figures from top to bottom represent birth weight, weaning weight, and yearling weight, respectively. The first column shows the *NCAPG-LCORL* locus as a QTL for growth trait (*26*). The color of each point represents the LD (r²) between each variant and rs384548488. The second column demonstrates a significant positive correlation between the -log10(*p* GWAS) of variants in average daily gain GWAS and their LD (R²) with rs384548488, suggesting that rs384548488 may be linked to growth traits. The red line is the nominal significance at *p* = 1 x 10^-5^.

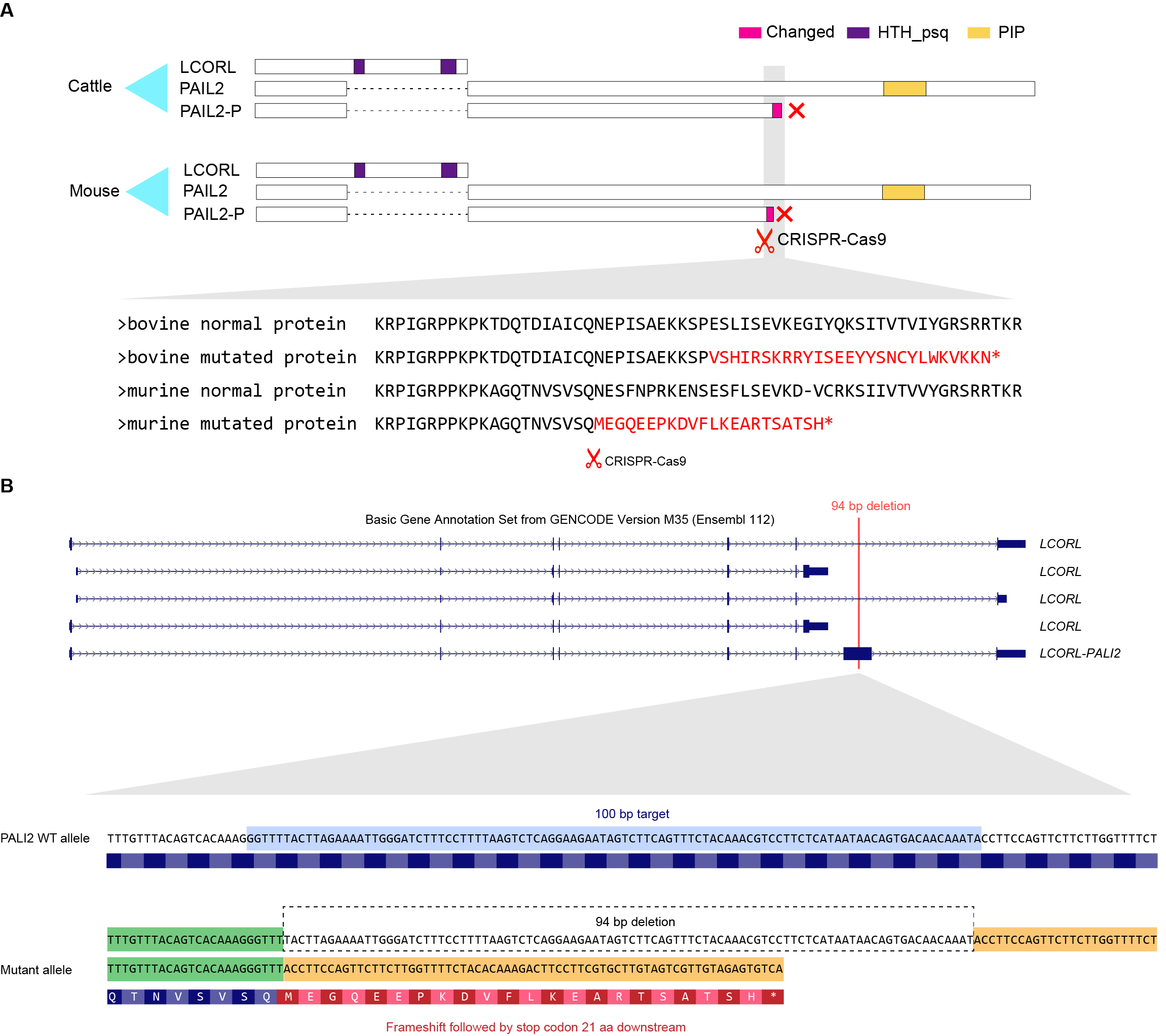

**Fig. S5.**

**The CRISPR targeting strategy for PALI2 in C57BL/6J mice.** **(A)** Top: A schematic view of the mouse LCORL and PALI2 protein isoforms. Bottom: Alignment of PALI2 protein sequences between cattle and mice. Bovine mutated protein sequence highlights the 27 amino acid substitution in cattle’s PALI2 protein caused by rs384548488, leading to a premature stop codon. Murine mutated protein sequence highlights a mutation introduced via CRISPR-Cas9 in mice, resulting in a 22 amino acid substitution and a premature stop codon in mouse PALI2 protein. **(B)** Top: A schematic view of the mouse LCORL and PALI2 transcript isoforms. Bottom: A guide RNA was designed to target a 100 bp region in the PALI2-specific exon (exon 8), which contains the PRC2 interaction domain. In the mutant allele, a 94 bp deletion was introduced, leading to premature translation termination of PALI2.

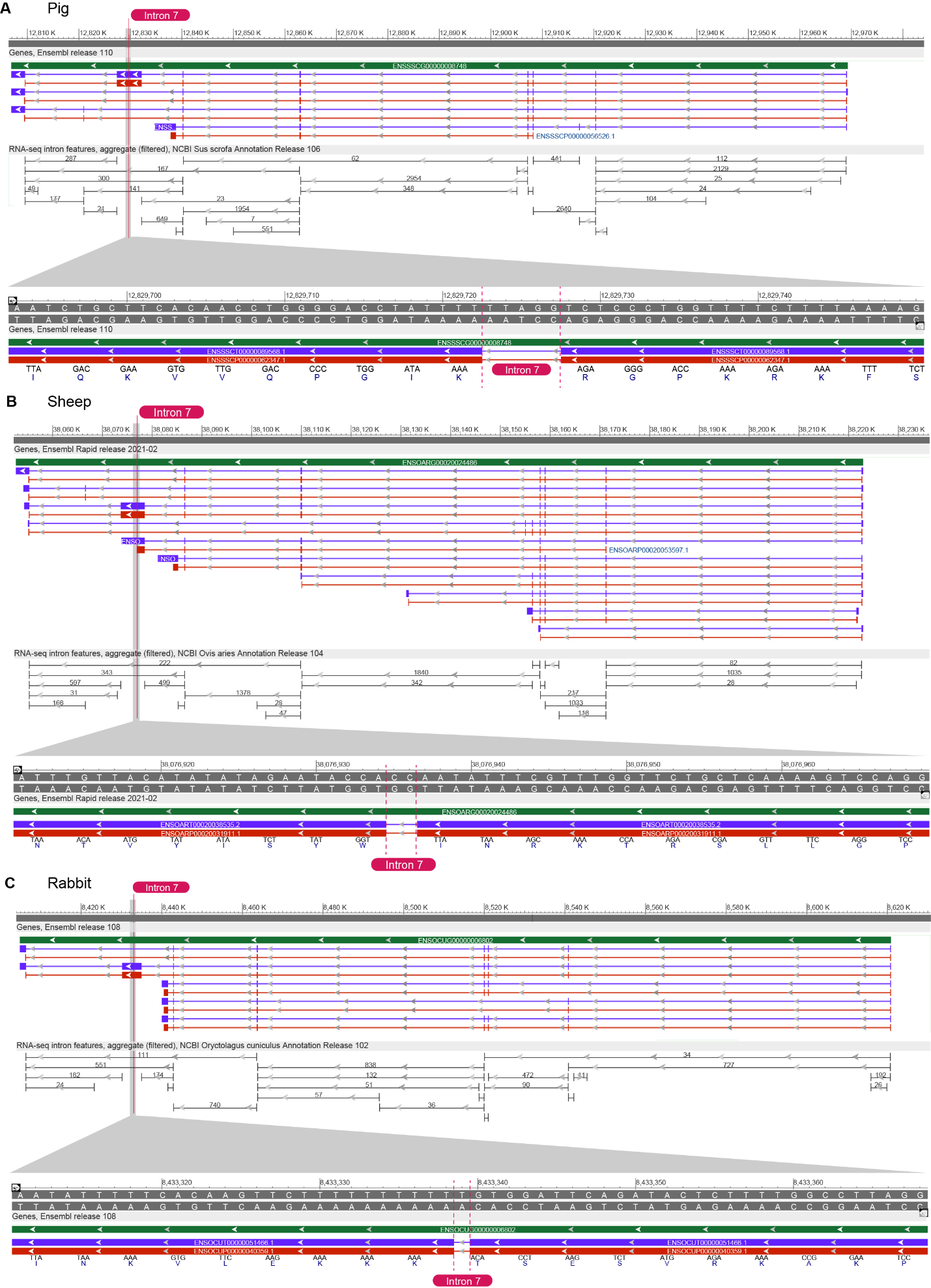

**Fig. S6.**

**Gene structure of the *LCORL* in Pig (A), Sheep (B), and Rabbit (C).** The first track shows the gene annotations provided by Ensembl. The second track displays the RNA-seq intron features from NCBI. The third track provides a close-up view of the frameshift intron. The frameshift intron (intron 7) is marked with red vertical lines. The lengths of these frameshift introns are all less than 5 bp, and no RNA-seq intron features are detected. Therefore, we speculate that these frameshift introns are not true introns but rather result from incorrect annotations due to the presence of pLoPD mutations in the reference genome.

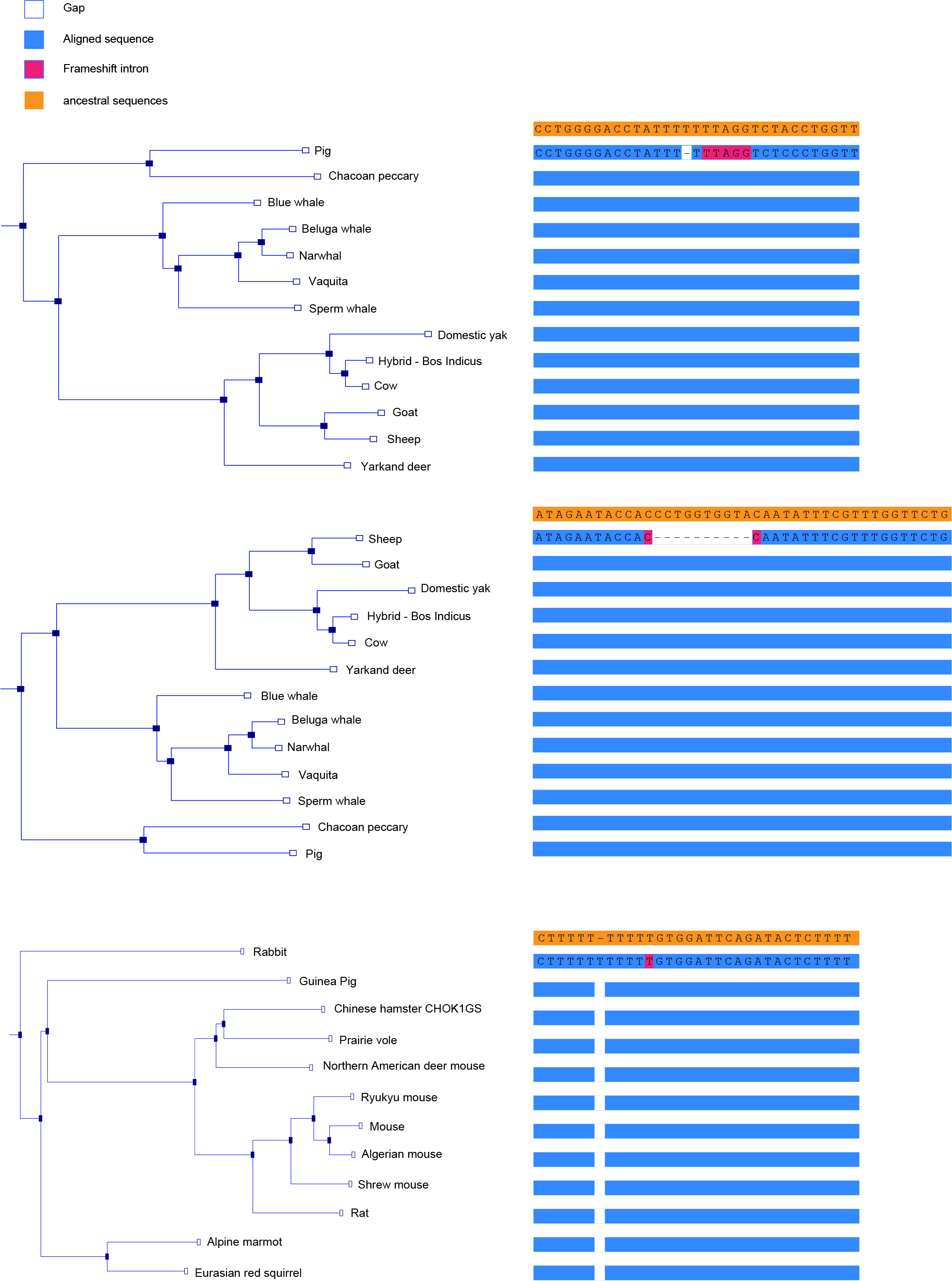

**Fig. S7.**

**Local multi-species alignments of the *LCORL* frameshift intron (intron 7) in Pig, Sheep, and Rabbit.** The multi-species sequence alignments and ancestral sequences were downloaded from Ensembl 43 eutherian mammals EPO dataset. The frameshift introns predicted by Ensembl are highlighted in red. Near each frameshift intron, there is a mutation that may explain the emergence of the frameshift intron.

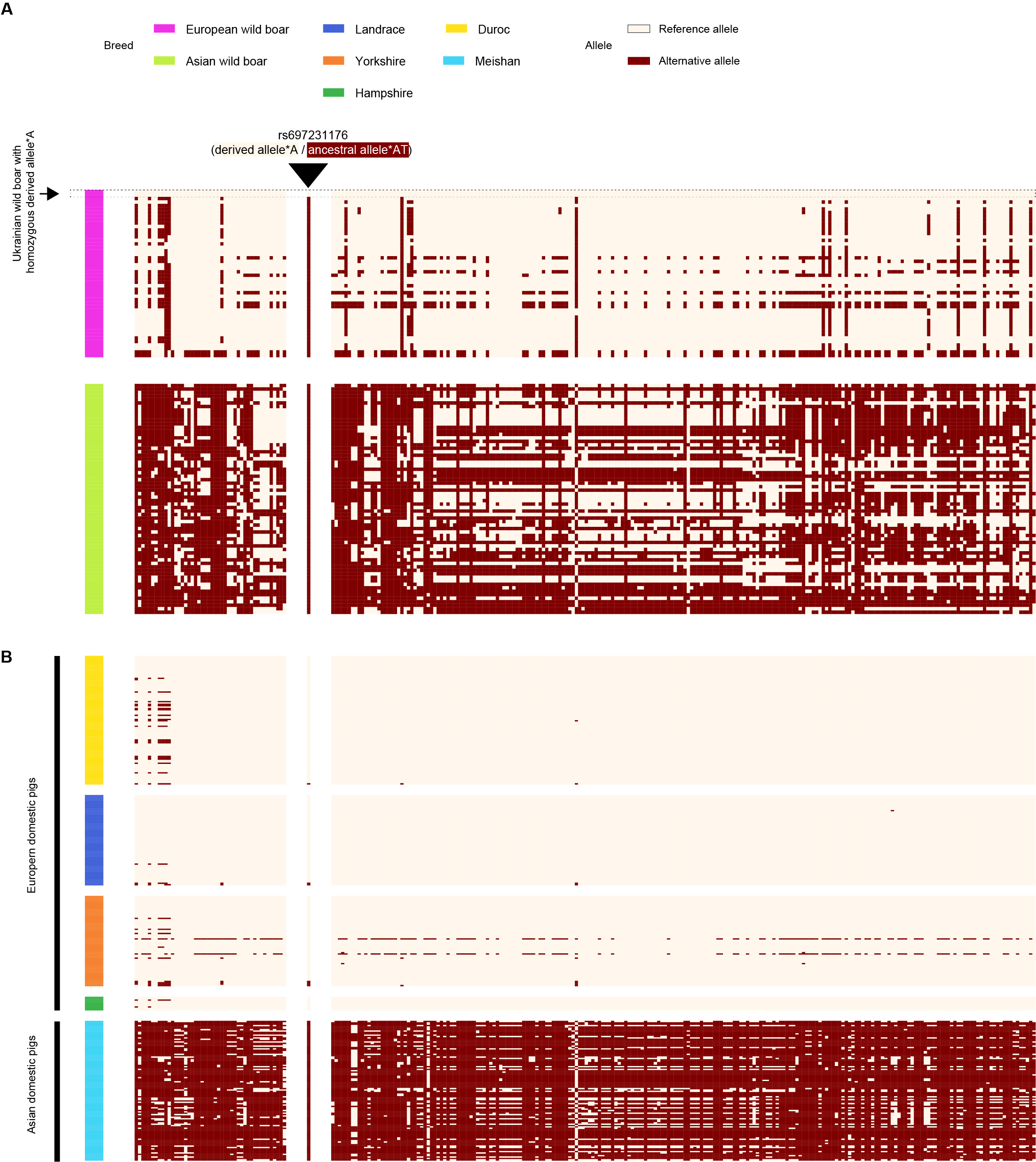

**Fig. S8.**

**Haplotype patterns at the *LCORL* locus in pigs.** **(A)** Haplotype patterns at the *LCORL* locus in European and Asian wild boars. Only one Ukrainian wild boar is homozygous for the derived allele (A) at rs697231176, while all other European and Asian wild boars carry the ancestral allele (AT) at rs697231176. **(B)** The haplotype pattern at the *LCORL* locus in European domesticated pigs shows signatures of selection. The derived allele (A) at rs697231176 is nearly fixed in European meat pigs. The pig breeds shown here are consistent with those used by Rubin et al. in their Sweep Analysis. Each row represents a haplotype, and each column represents a locus. The positions along the x-axis are shared between panels A and B.

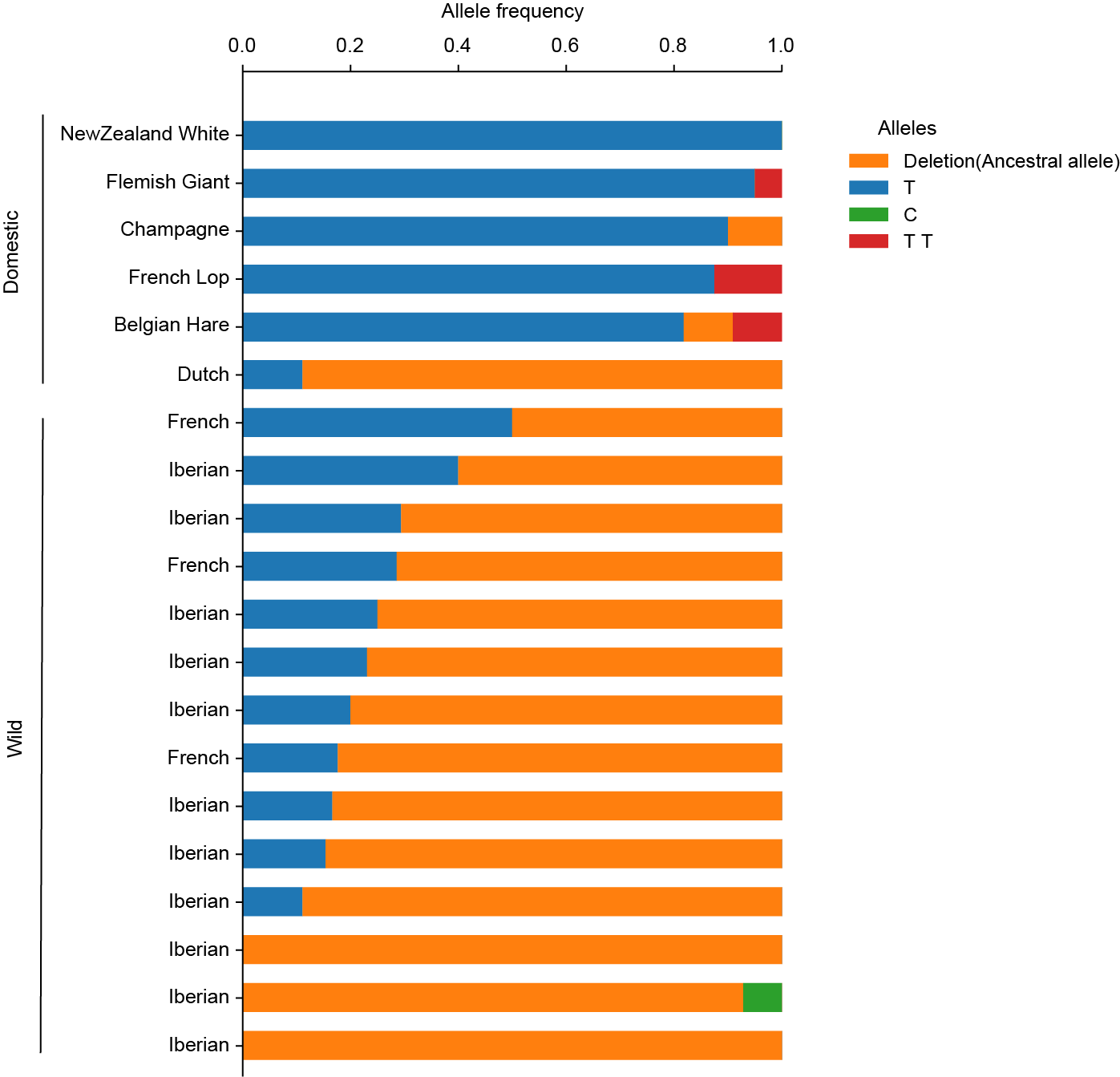

**Fig. S9.**

**Allele frequencies at the Chr2:8433329 locus in domestic and wild rabbit populations.** Allele frequencies were calculated from pooled whole-genome resequencing data published by M. Carneiro et al. The populations presented align with those used in M. Carneiro et al.'s study. The x-axis shows allele frequencies inferred from reads count, and the y-axis shows the corresponding rabbit populations. Sample information is available in Table S7.

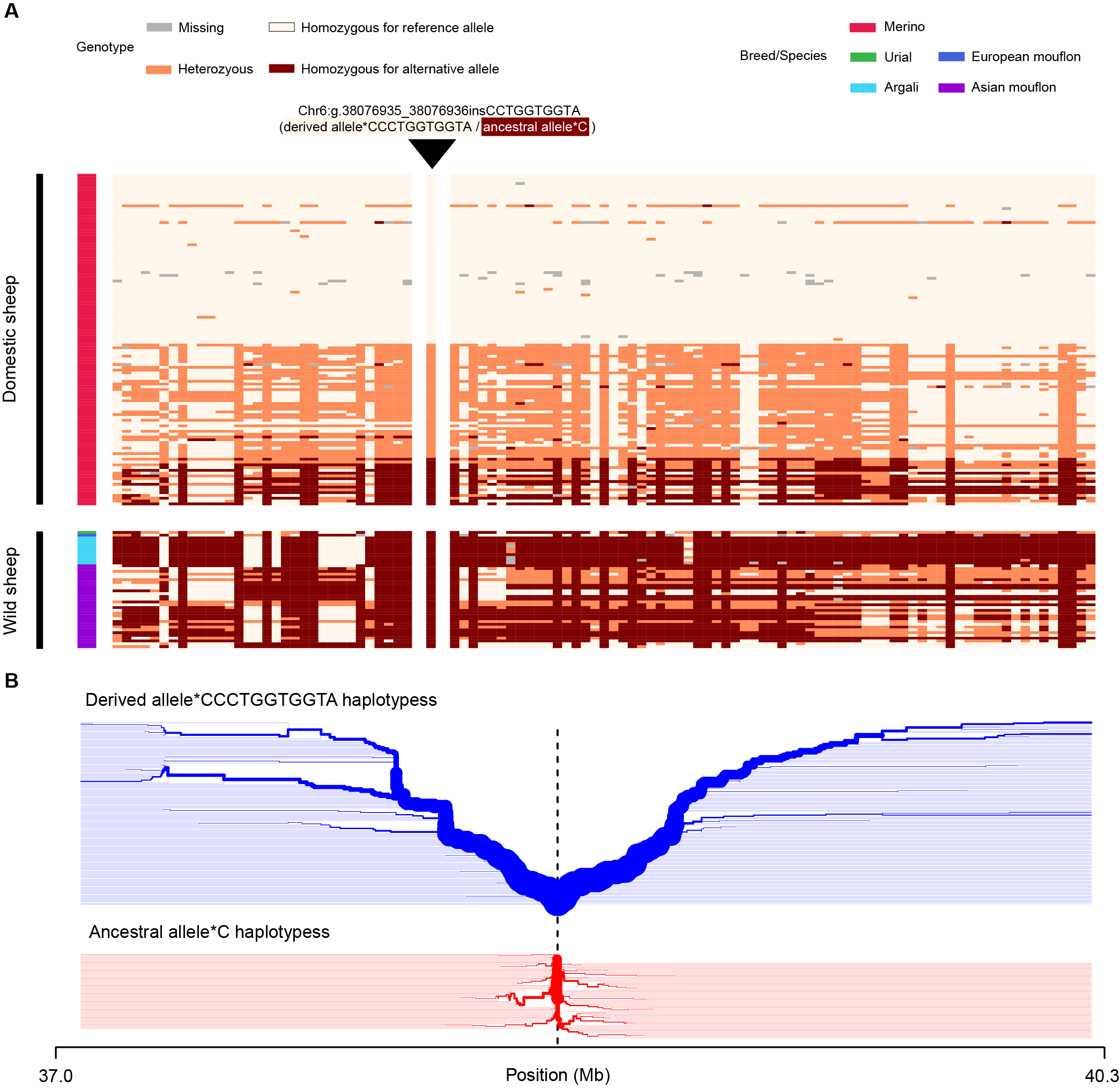

**Fig. S10.**

**The genotype and haplotype patterns at the *LCORL* locus in sheep. (A)** The genotype patterns at the *LCORL* region (Chr6:38052622-38222929) in sheep. The pLoPD mutation Chr6:g.38076935_38076936insCCTGGTGGTA is polymorphic in domestic sheep but absent in wild sheep. **(B)** Bifurcation diagram for haplotypes in Merino sheep, starting from the pLoPD mutation Chr6:g.38076935_38076936insCCTGGTGGTA. In Merino sheep, haplotypes carrying the pLoPD mutations exhibit longer extended haplotype than haplotypes without the pLoPD mutations, suggesting that haplotypes carrying the pLoPD mutations may be under positive selection.

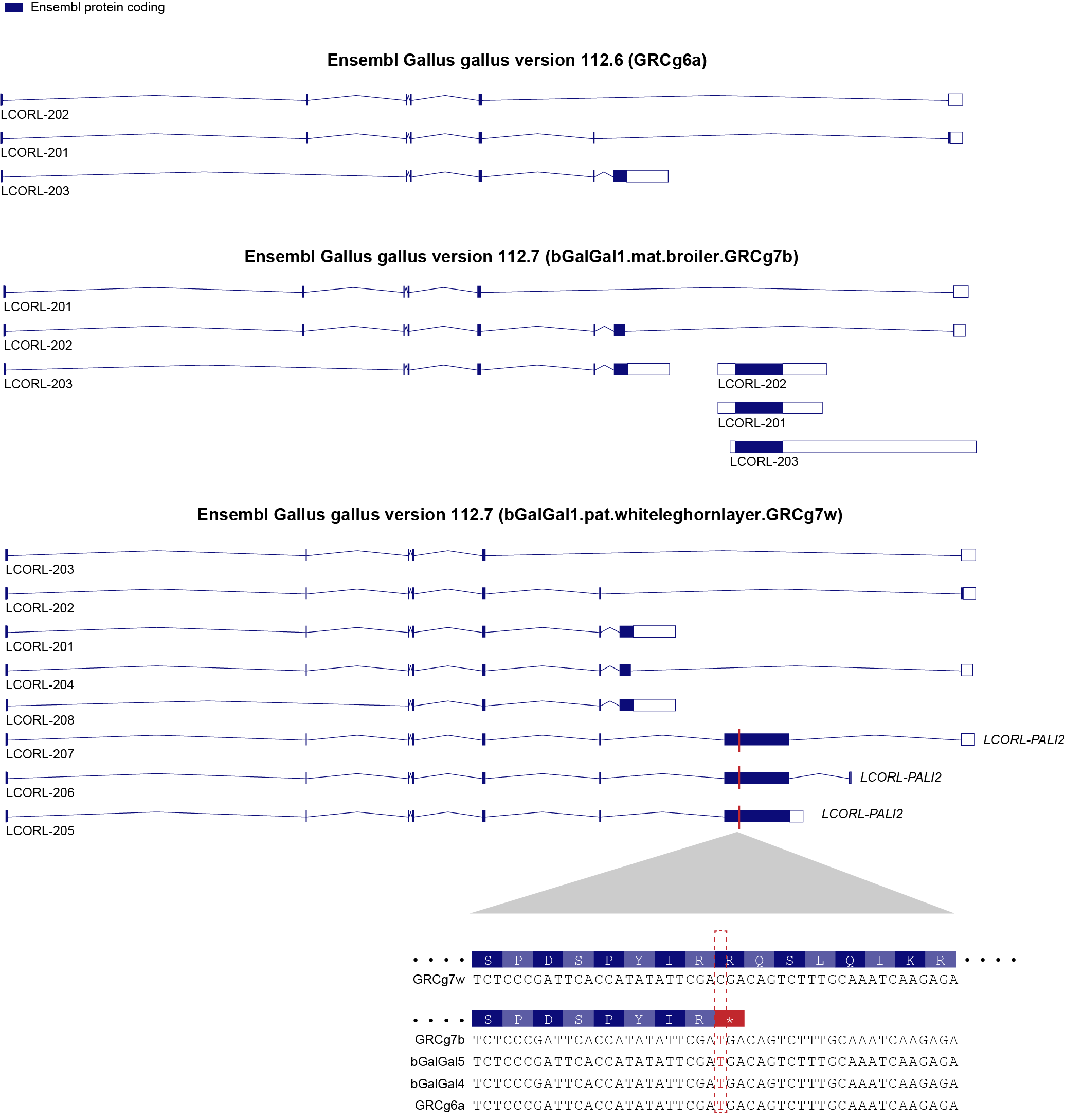

**Fig. S11.**

**Predicted gene structure of *LCORL* in different chicken reference genomes.** The PALI2 isoform is not annotated in the reference genomes GRCg6a and GRCg7b, but it is annotated in the reference genome GRCg7w. The sequences flanking the pLoPD variant rs317817652 are shown at the bottom. The C-to-G nonsense mutation occurs at rs317817652 in the reference genomes GRCg6a, GRCg7b, as well as in genomes bGalGal5b and GalGal4. The pLoPD variant rs317817652 in domesticated chickens is marked by a red vertical line.

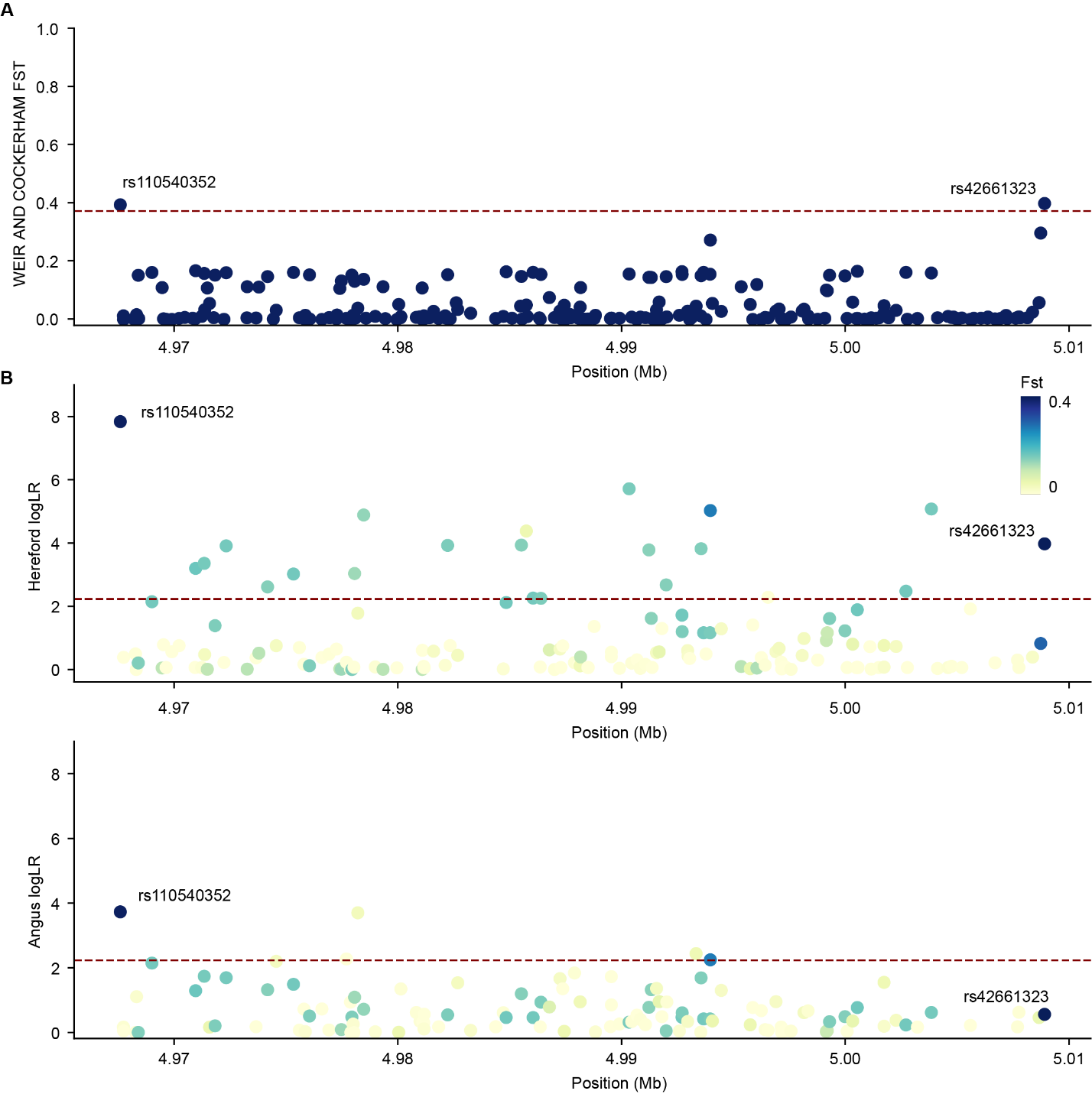

**Fig. S12.**

**Selective Sweep Analysis at the *STC2* Locus.** **(A)** In the body size QTL region Chr20:4967596-5008911, *Fst* values were calculated between British and Continental European cattle. The red line represents the genome-wide significance threshold at Fst = 0.37. Only two variants, rs110540352 and rs42661323, exhibit strong differentiation. **(B)** The CLUES analysis was performed for all biallelic SNVs in the body size QTL region Chr20:4967596-5008911, comparing Hereford and Angus cattle. The red line indicates the significance threshold at logLR = 2.23. The variant rs110540352 is identified as the lead SNV in both Hereford and Angus populations.

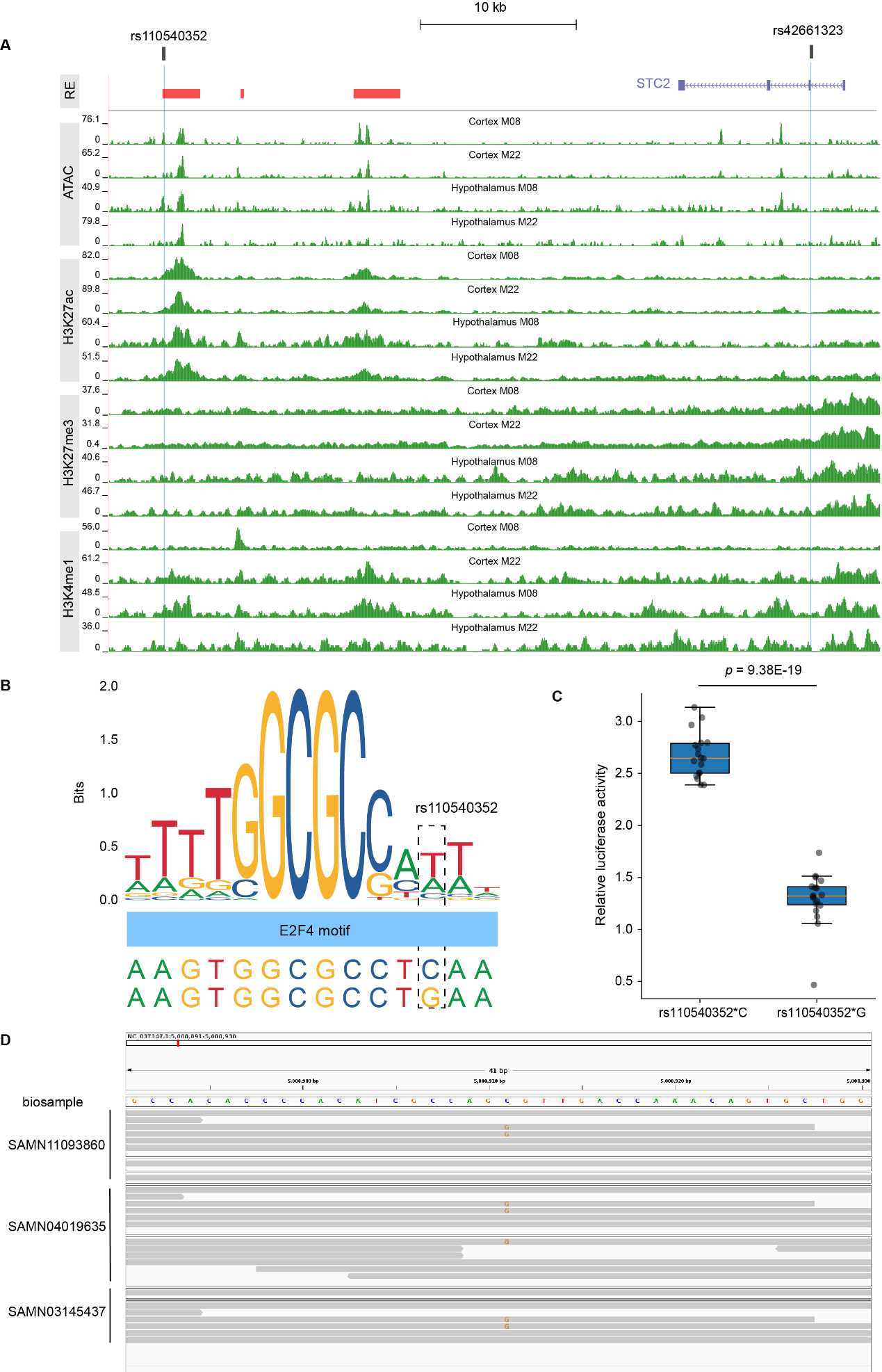

**Fig. S13.**

**Regulatory variant rs110540352 impacts STC2 expression.** **(A)** rs110540352 is located in an active regulatory element in the cerebral cortex and hypothalamus. Red rectangles are active regulatory elements in the cerebral cortex and hypothalamus. Green tracks represent ATAC, H3K27ac, H3K27me3, and H3K4me1 data in the cerebral cortex and hypothalamus. **(B)** rs110540352 overlaps transcription factor binding sites. The C-to-G mutation at rs110540352 disrupts the binding motif of E2F4. **(C)** Dual-luciferase assay results demonstrating that the C-to-G mutation at rs110540352 decreases luciferase activity. The *P-value* was calculated using a two-sided Student’s t-test. **(D)** isualizing RNA-seq reads of the cortex from rs42661323 heterozygous Herefords. Biosample SAMN11093860 contains tracks from the Temporal cortex, Frontal cortex, and Cerebral cortex, from top to bottom Biosample SAMN04019635 includes two RNA-seq tracks from the Cerebral cortex. Biosample SAMN03145437 contains tracks from the Frontal cortex and Cerebral cortex, from top to bottom. Due to the low number of reads covering rs42661323 (20:5008911) in each sample, we combined the samples from the cerebral cortex. There are fewer reads carrying the derived allele rs42661323*G (n=7) than the ancestral allele rs42661323*C (n=16) (Binomial one-sided test, *p*=0.047).

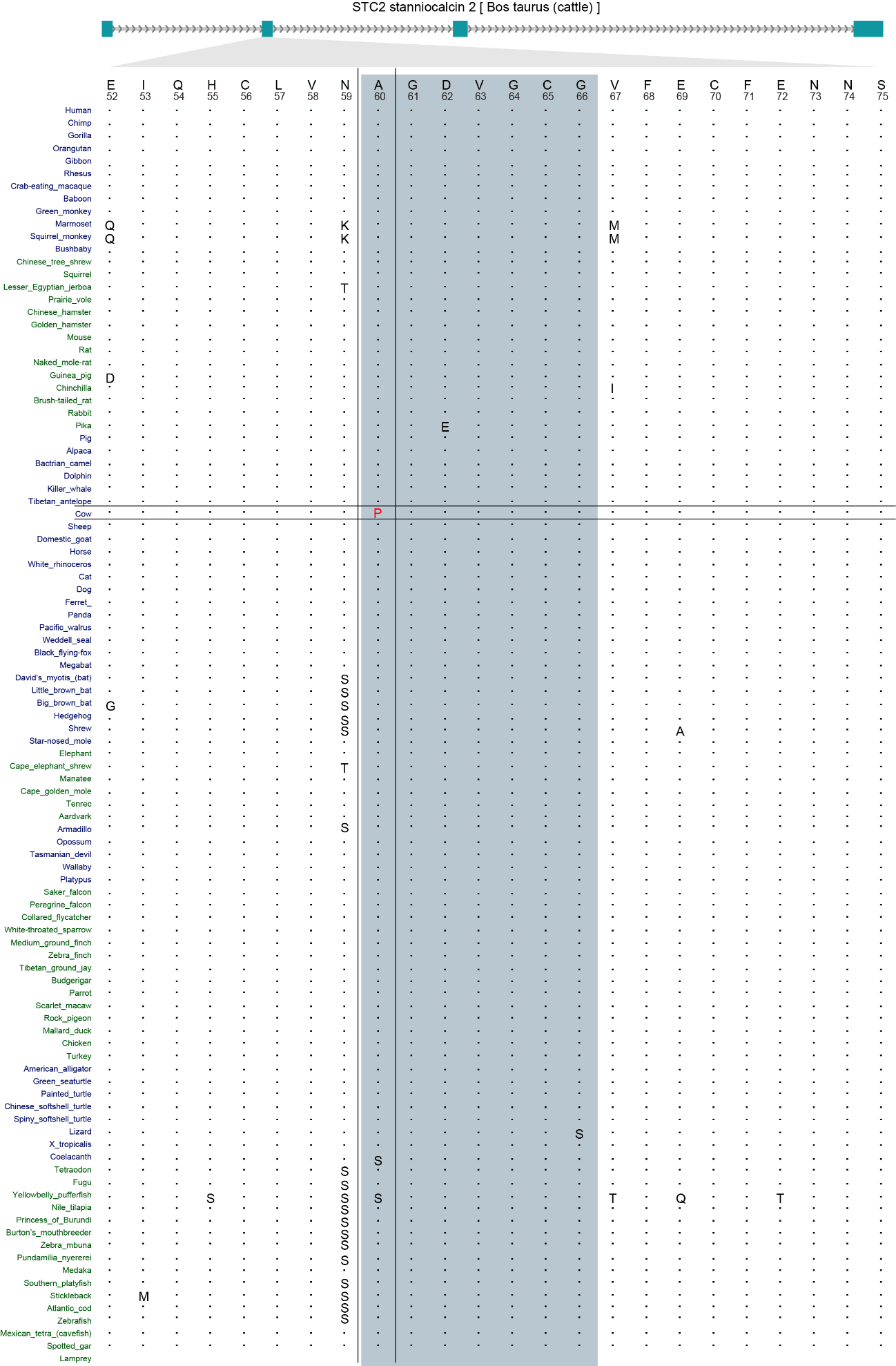

**Fig. S14.**

**Conservation analysis of the 60th amino acid of STC2.** The 60-66th amino acids of STC2 is the interaction region with PAPP-A, marked with blue shading. Sequence alignments were obtained from UCSC Vertebrate Multiz Alignment & Conservation (100 Species).

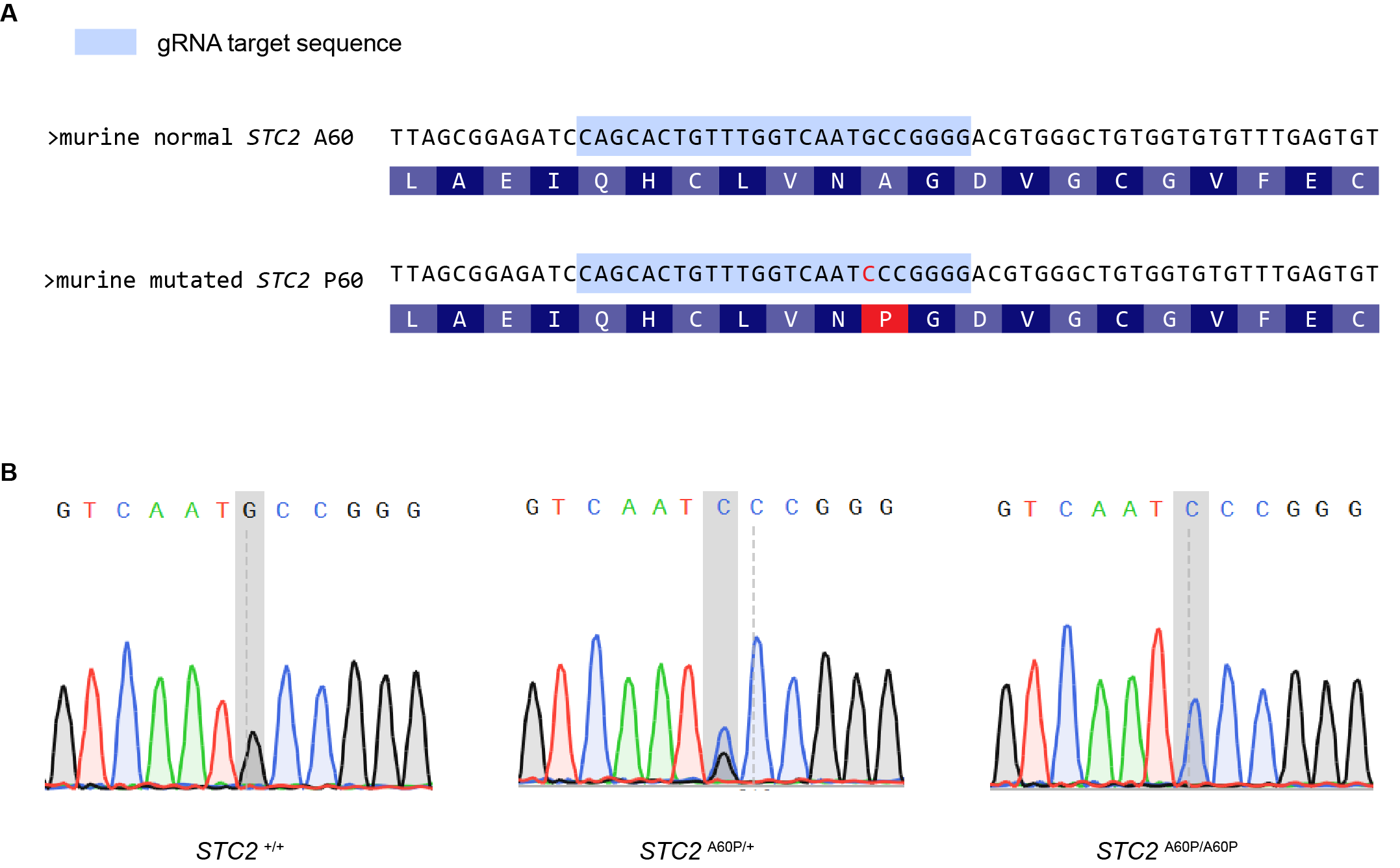
**Fig. S15.**

**Generation of STC2 A60P mutation mice.** **(A)** The schematic diagram of STC2 A60P generation by CRISPR/Cas9. The edited base is indicated in red. **(B)** Sanger sequencing showed sequencing peaks in *STC2*^+/+^, *STC2*^A60P/+^, *STC2*^A60P/A60P^, respectively.

**Table S6.**

**The domestic pigs and wild boars used in this study.** Sequencing depth is given relative to the pig reference genome.

| **Run ID** | **Domestic/Wild** | **Population** | **Group Fig. S7** | **Depth** | **BioProject** |
| --- | --- | --- | --- | --- | --- |
| ERS177302 | Domestic | Duroc | Duroc | 5.5 | PRJEB1683 |
| ERS177303 | Domestic | Duroc | Duroc | 6.2 | PRJEB1683 |
| ERS177305 | Domestic | Duroc | Duroc | 5.1 | PRJEB1683 |
| ERS804986 | Domestic | Duroc | Duroc | 11.8 | PRJEB9922 |
| ERS804987 | Domestic | Duroc | Duroc | 11.3 | PRJEB9922 |
| ERS804988 | Domestic | Duroc | Duroc | 5.1 | PRJEB9922 |
| ERS804989 | Domestic | Duroc | Duroc | 6.4 | PRJEB9922 |
| SRS562607 | Domestic | Duroc | Duroc | 9.3 | PRJNA239399 |
| SRS703257 | Domestic | Duroc | Duroc | 5.7 | PRJNA260763 |
| SRS703258 | Domestic | Duroc | Duroc | 12.3 | PRJNA260763 |
| SRS703262 | Domestic | Duroc | Duroc | 11.1 | PRJNA260763 |
| SRS703263 | Domestic | Duroc | Duroc | 11.0 | PRJNA260763 |
| SRS703264 | Domestic | Duroc | Duroc | 11.4 | PRJNA260763 |
| SRS703265 | Domestic | Duroc | Duroc | 10.3 | PRJNA260763 |
| SRS703266 | Domestic | Duroc | Duroc | 10.3 | PRJNA260763 |
| SRS703267 | Domestic | Duroc | Duroc | 11.7 | PRJNA260763 |
| SRS703271 | Domestic | Duroc | Duroc | 10.2 | PRJNA260763 |
| SRS703272 | Domestic | Duroc | Duroc | 10.4 | PRJNA260763 |
| SRS703273 | Domestic | Duroc | Duroc | 10.7 | PRJNA260763 |
| SRS703274 | Domestic | Duroc | Duroc | 11.3 | PRJNA260763 |
| SRS703275 | Domestic | Duroc | Duroc | 10.5 | PRJNA260763 |
| SRS703276 | Domestic | Duroc | Duroc | 11.0 | PRJNA260763 |
| SRS703277 | Domestic | Duroc | Duroc | 10.2 | PRJNA260763 |
| SRS703278 | Domestic | Duroc | Duroc | 10.9 | PRJNA260763 |
| SRS703279 | Domestic | Duroc | Duroc | 10.7 | PRJNA260763 |
| SRS703259 | Domestic | Duroc | Duroc | 11.6 | PRJNA260763 |
| SRS1725712 | Domestic | Duroc | Duroc | 11.8 | PRJNA343658 |
| SRS1725715 | Domestic | Duroc | Duroc | 10.5 | PRJNA343658 |
| SRS1725716 | Domestic | Duroc | Duroc | 13.6 | PRJNA343658 |
| SRS1725717 | Domestic | Duroc | Duroc | 17.1 | PRJNA343658 |
| SRS1725718 | Domestic | Duroc | Duroc | 19.3 | PRJNA343658 |
| SRS1725719 | Domestic | Duroc | Duroc | 11.2 | PRJNA343658 |
| SRS1725721 | Domestic | Duroc | Duroc | 12.2 | PRJNA343658 |
| SRS1725722 | Domestic | Duroc | Duroc | 8.9 | PRJNA343658 |
| SRS1725723 | Domestic | Duroc | Duroc | 7.7 | PRJNA343658 |
| SRS1725724 | Domestic | Duroc | Duroc | 11.7 | PRJNA343658 |
| SRS2043289 | Domestic | Duroc | Duroc | 11.2 | PRJNA378496 |
| SRS2043844 | Domestic | Duroc | Duroc | 10.3 | PRJNA378496 |
| SRS2043851 | Domestic | Duroc | Duroc | 5.1 | PRJNA378496 |
| SRS2043855 | Domestic | Duroc | Duroc | 10.3 | PRJNA378496 |
| SRS2043859 | Domestic | Duroc | Duroc | 9.6 | PRJNA378496 |
| SRS2043983 | Domestic | Duroc | Duroc | 9.6 | PRJNA378496 |
| SRS2043998 | Domestic | Duroc | Duroc | 6.6 | PRJNA378496 |
| SRS2045986 | Domestic | Duroc | Duroc | 8.6 | PRJNA378496 |
| SRS2051177 | Domestic | Duroc | Duroc | 6.4 | PRJNA378496 |
| SRS2053398 | Domestic | Duroc | Duroc | 5.6 | PRJNA378496 |
| SRS2058801 | Domestic | Duroc | Duroc | 8.8 | PRJNA378496 |
| SRS4099336 | Domestic | Duroc | Duroc | 26.1 | PRJNA507853 |
| ERS177306 | Domestic | Hampshire | Hampshire | 6.2 | PRJEB1683 |
| ERS177307 | Domestic | Hampshire | Hampshire | 5.8 | PRJEB1683 |
| ERS804992 | Domestic | Hampshire | Hampshire | 8.5 | PRJEB9922 |
| ERS804993 | Domestic | Hampshire | Hampshire | 8.1 | PRJEB9922 |
| SRS1261795 | Domestic | Hampshire | Hampshire | 58.5 | PRJNA309108 |
| ERS177314 | Domestic | Landrace | Landrace | 5.3 | PRJEB1683 |
| ERS177312 | Domestic | Landrace | Landrace | 6.4 | PRJEB1683 |
| ERS177313 | Domestic | Landrace | Landrace | 7.3 | PRJEB1683 |
| ERS804996 | Domestic | Landrace | Landrace | 7.8 | PRJEB9922 |
| ERS804997 | Domestic | Landrace | Landrace | 14.4 | PRJEB9922 |
| ERS804999 | Domestic | Landrace | Landrace | 6.6 | PRJEB9922 |
| ERS805000 | Domestic | Landrace | Landrace | 9.0 | PRJEB9922 |
| SRS703281 | Domestic | Landrace | Landrace | 10.2 | PRJNA260763 |
| SRS703282 | Domestic | Landrace | Landrace | 10.3 | PRJNA260763 |
| SRS703283 | Domestic | Landrace | Landrace | 9.8 | PRJNA260763 |
| SRS703284 | Domestic | Landrace | Landrace | 9.5 | PRJNA260763 |
| SRS703285 | Domestic | Landrace | Landrace | 6.8 | PRJNA260763 |
| SRS703286 | Domestic | Landrace | Landrace | 6.3 | PRJNA260763 |
| SRS703287 | Domestic | Landrace | Landrace | 5.8 | PRJNA260763 |
| SRS703289 | Domestic | Landrace | Landrace | 8.3 | PRJNA260763 |
| SRS703290 | Domestic | Landrace | Landrace | 8.0 | PRJNA260763 |
| SRS703291 | Domestic | Landrace | Landrace | 9.5 | PRJNA260763 |
| SRS703292 | Domestic | Landrace | Landrace | 9.4 | PRJNA260763 |
| SRS703293 | Domestic | Landrace | Landrace | 9.8 | PRJNA260763 |
| SRS1261807 | Domestic | Landrace | Landrace | 52.9 | PRJNA309108 |
| SRS1725708 | Domestic | Landrace | Landrace | 16.8 | PRJNA343658 |
| SRS1725709 | Domestic | Landrace | Landrace | 15.9 | PRJNA343658 |
| SRS1725710 | Domestic | Landrace | Landrace | 10.8 | PRJNA343658 |
| SRS1725713 | Domestic | Landrace | Landrace | 13.4 | PRJNA343658 |
| SRS1725720 | Domestic | Landrace | Landrace | 11.0 | PRJNA343658 |
| SRS1725731 | Domestic | Landrace | Landrace | 12.6 | PRJNA343658 |
| SRS1725743 | Domestic | Landrace | Landrace | 17.6 | PRJNA343658 |
| SRS1725753 | Domestic | Landrace | Landrace | 18.3 | PRJNA343658 |
| SRS1725764 | Domestic | Landrace | Landrace | 10.4 | PRJNA343658 |
| SRS1725775 | Domestic | Landrace | Landrace | 13.6 | PRJNA343658 |
| SRS1725778 | Domestic | Landrace | Landrace | 17.8 | PRJNA343658 |
| SRS2043847 | Domestic | Landrace | Landrace | 9.6 | PRJNA378496 |
| SRS4099337 | Domestic | Landrace | Landrace | 31.8 | PRJNA507853 |
| ERS485866 | Domestic | Meishan | Meishan | 7.2 | PRJEB1683 |
| ERS485867 | Domestic | Meishan | Meishan | 7.4 | PRJEB1683 |
| ERS485868 | Domestic | Meishan | Meishan | 5.3 | PRJEB1683 |
| ERS485869 | Domestic | Meishan | Meishan | 6.1 | PRJEB1683 |
| ERS485870 | Domestic | Meishan | Meishan | 7.5 | PRJEB1683 |
| ERS485871 | Domestic | Meishan | Meishan | 6.3 | PRJEB1683 |
| ERS497905 | Domestic | Meishan | Meishan | 7.2 | PRJEB1683 |
| ERS497906 | Domestic | Meishan | Meishan | 7.4 | PRJEB1683 |
| ERS177331 | Domestic | Meishan | Meishan | 6.2 | PRJEB1683 |
| ERS177332 | Domestic | Meishan | Meishan | 6.1 | PRJEB1683 |
| ERS177333 | Domestic | Meishan | Meishan | 5.6 | PRJEB1683 |
| ERS177334 | Domestic | Meishan | Meishan | 6.9 | PRJEB1683 |
| ERS804949 | Domestic | Meishan | Meishan | 9.4 | PRJEB9922 |
| ERS804950 | Domestic | Meishan | Meishan | 9.9 | PRJEB9922 |
| ERS804951 | Domestic | Meishan | Meishan | 8.0 | PRJEB9922 |
| ERS804952 | Domestic | Meishan | Meishan | 7.9 | PRJEB9922 |
| ERS804953 | Domestic | Meishan | Meishan | 7.0 | PRJEB9922 |
| ERS804954 | Domestic | Meishan | Meishan | 8.1 | PRJEB9922 |
| ERS804955 | Domestic | Meishan | Meishan | 10.1 | PRJEB9922 |
| ERS804956 | Domestic | Meishan | Meishan | 8.4 | PRJEB9922 |
| ERS804957 | Domestic | Meishan | Meishan | 8.5 | PRJEB9922 |
| ERS804958 | Domestic | Meishan | Meishan | 12.1 | PRJEB9922 |
| SRS1261813 | Domestic | Meishan | Meishan | 61.6 | PRJNA309108 |
| SRS2040260 | Domestic | Meishan | Meishan | 7.9 | PRJNA378496 |
| SRS2040900 | Domestic | Meishan | Meishan | 7.6 | PRJNA378496 |
| SRS2043292 | Domestic | Meishan | Meishan | 9.9 | PRJNA378496 |
| SRS2043301 | Domestic | Meishan | Meishan | 8.5 | PRJNA378496 |
| SRS2043302 | Domestic | Meishan | Meishan | 10.4 | PRJNA378496 |
| SRS2043609 | Domestic | Meishan | Meishan | 8.9 | PRJNA378496 |
| SRS2043845 | Domestic | Meishan | Meishan | 5.1 | PRJNA378496 |
| SRS2043849 | Domestic | Meishan | Meishan | 9.6 | PRJNA378496 |
| SRS2043852 | Domestic | Meishan | Meishan | 8.6 | PRJNA378496 |
| SRS2043854 | Domestic | Meishan | Meishan | 9.1 | PRJNA378496 |
| SRS2043856 | Domestic | Meishan | Meishan | 9.2 | PRJNA378496 |
| SRS2043915 | Domestic | Meishan | Meishan | 10.1 | PRJNA378496 |
| SRS2043995 | Domestic | Meishan | Meishan | 9.1 | PRJNA378496 |
| SRS2044030 | Domestic | Meishan | Meishan | 10.6 | PRJNA378496 |
| SRS2044055 | Domestic | Meishan | Meishan | 7.4 | PRJNA378496 |
| SRS2044542 | Domestic | Meishan | Meishan | 8.6 | PRJNA378496 |
| SRS2046192 | Domestic | Meishan | Meishan | 8.0 | PRJNA378496 |
| SRS2046488 | Domestic | Meishan | Meishan | 8.8 | PRJNA378496 |
| SRS2048171 | Domestic | Meishan | Meishan | 8.2 | PRJNA378496 |
| SRS2048172 | Domestic | Meishan | Meishan | 8.7 | PRJNA378496 |
| SRS2048245 | Domestic | Meishan | Meishan | 9.4 | PRJNA378496 |
| SRS2048284 | Domestic | Meishan | Meishan | 9.1 | PRJNA378496 |
| SRS2048767 | Domestic | Meishan | Meishan | 5.2 | PRJNA378496 |
| SRS2048848 | Domestic | Meishan | Meishan | 9.5 | PRJNA378496 |
| SRS2051325 | Domestic | Meishan | Meishan | 6.6 | PRJNA378496 |
| SRS2052083 | Domestic | Meishan | Meishan | 8.9 | PRJNA378496 |
| SRS2053229 | Domestic | Meishan | Meishan | 8.3 | PRJNA378496 |
| SRS2053343 | Domestic | Meishan | Meishan | 10.2 | PRJNA378496 |
| ERS177318 | Domestic | Yorkshire | Yorkshire | 6.8 | PRJEB1683 |
| ERS177319 | Domestic | Yorkshire | Yorkshire | 6.8 | PRJEB1683 |
| ERS177320 | Domestic | Yorkshire | Yorkshire | 6.9 | PRJEB1683 |
| ERS177321 | Domestic | Yorkshire | Yorkshire | 6.4 | PRJEB1683 |
| ERS177322 | Domestic | Yorkshire | Yorkshire | 5.9 | PRJEB1683 |
| ERS177323 | Domestic | Yorkshire | Yorkshire | 6.6 | PRJEB1683 |
| ERS177325 | Domestic | Yorkshire | Yorkshire | 6.9 | PRJEB1683 |
| ERS177326 | Domestic | Yorkshire | Yorkshire | 5.5 | PRJEB1683 |
| ERS177327 | Domestic | Yorkshire | Yorkshire | 5.6 | PRJEB1683 |
| ERS177328 | Domestic | Yorkshire | Yorkshire | 6.1 | PRJEB1683 |
| ERS177329 | Domestic | Yorkshire | Yorkshire | 5.6 | PRJEB1683 |
| ERS177330 | Domestic | Yorkshire | Yorkshire | 5.6 | PRJEB1683 |
| ERS804938 | Domestic | Yorkshire | Yorkshire | 7.2 | PRJEB9922 |
| ERS804939 | Domestic | Yorkshire | Yorkshire | 8.8 | PRJEB9922 |
| ERS805002 | Domestic | Yorkshire | Yorkshire | 8.1 | PRJEB9922 |
| SRS2168992 | Domestic | Yorkshire | Yorkshire | 10.7 | PRJNA255085 |
| SRS703317 | Domestic | Yorkshire | Yorkshire | 8.3 | PRJNA260763 |
| SRS703319 | Domestic | Yorkshire | Yorkshire | 8.7 | PRJNA260763 |
| SRS703320 | Domestic | Yorkshire | Yorkshire | 8.0 | PRJNA260763 |
| SRS703321 | Domestic | Yorkshire | Yorkshire | 7.4 | PRJNA260763 |
| SRS703322 | Domestic | Yorkshire | Yorkshire | 7.1 | PRJNA260763 |
| SRS703323 | Domestic | Yorkshire | Yorkshire | 8.6 | PRJNA260763 |
| SRS703324 | Domestic | Yorkshire | Yorkshire | 8.6 | PRJNA260763 |
| SRS703325 | Domestic | Yorkshire | Yorkshire | 6.1 | PRJNA260763 |
| SRS703326 | Domestic | Yorkshire | Yorkshire | 6.5 | PRJNA260763 |
| SRS703331 | Domestic | Yorkshire | Yorkshire | 10.0 | PRJNA260763 |
| SRS703332 | Domestic | Yorkshire | Yorkshire | 10.8 | PRJNA260763 |
| SRS703333 | Domestic | Yorkshire | Yorkshire | 10.6 | PRJNA260763 |
| SRS703334 | Domestic | Yorkshire | Yorkshire | 10.4 | PRJNA260763 |
| SRS703336 | Domestic | Yorkshire | Yorkshire | 10.4 | PRJNA260763 |
| SRS1261809 | Domestic | Yorkshire | Yorkshire | 55.2 | PRJNA309108 |
| SRS2043916 | Domestic | Yorkshire | Yorkshire | 9.9 | PRJNA378496 |
| SRS4099338 | Domestic | Yorkshire | Yorkshire | 26.7 | PRJNA507853 |
| ERS485880 | Wild | Chinese wild boar | Asian wild boar | 8.0 | PRJEB1683 |
| ERS177354 | Wild | Chinese wild boar | Asian wild boar | 6.7 | PRJEB1683 |
| ERS485878 | Wild | Chinese wild boar | Asian wild boar | 6.3 | PRJEB1683 |
| ERS485879 | Wild | Chinese wild boar | Asian wild boar | 9.0 | PRJEB1683 |
| ERS177352 | Wild | Chinese wild boar | Asian wild boar | 6.9 | PRJEB1683 |
| ERS804969 | Wild | Chinese wild boar | Asian wild boar | 5.1 | PRJEB9922 |
| ERS804970 | Wild | Chinese wild boar | Asian wild boar | 8.2 | PRJEB9922 |
| ERS804971 | Wild | Chinese wild boar | Asian wild boar | 11.2 | PRJEB9922 |
| ERS804964 | Wild | Chinese wild boar | Asian wild boar | 5.2 | PRJEB9922 |
| ERS804965 | Wild | Chinese wild boar | Asian wild boar | 8.6 | PRJEB9922 |
| ERS804967 | Wild | Chinese wild boar | Asian wild boar | 28.2 | PRJEB9922 |
| ERS804968 | Wild | Chinese wild boar | Asian wild boar | 12.0 | PRJEB9922 |
| SRS465716 | Wild | Chinese wild boar | Asian wild boar | 21.6 | PRJNA213179 |
| SRS465717 | Wild | Chinese wild boar | Asian wild boar | 23.6 | PRJNA213179 |
| SRS465718 | Wild | Chinese wild boar | Asian wild boar | 22.1 | PRJNA213179 |
| SRS465719 | Wild | Chinese wild boar | Asian wild boar | 17.1 | PRJNA213179 |
| SRS465720 | Wild | Chinese wild boar | Asian wild boar | 16.2 | PRJNA213179 |
| SRS465721 | Wild | Chinese wild boar | Asian wild boar | 17.2 | PRJNA213179 |
| SRS1232269 | Wild | Chinese wild boar | Asian wild boar | 6.5 | PRJNA305081 |
| SRS1232279 | Wild | Chinese wild boar | Asian wild boar | 10.1 | PRJNA305081 |
| SRS1232297 | Wild | Chinese wild boar | Asian wild boar | 10.3 | PRJNA305081 |
| SRS2043996 | Wild | Chinese wild boar | Asian wild boar | 13.7 | PRJNA378496 |
| ERS177349 | Wild | French wild boar | European wild boar | 6.8 | PRJEB1683 |
| ERS805025 | Wild | French wild boar | European wild boar | 8.5 | PRJEB9922 |
| ERS805029 | Wild | Greek wild boar | European wild boar | 5.2 | PRJEB9922 |
| ERS805027 | Wild | Italian wild boar | European wild boar | 12.1 | PRJEB9922 |
| ERS805028 | Wild | Italian wild boar | European wild boar | 10.4 | PRJEB9922 |
| ERS805035 | Wild | Italian wild boar | European wild boar | 11.6 | PRJEB9922 |
| ERS805036 | Wild | Italian wild boar | European wild boar | 11.1 | PRJEB9922 |
| ERS805037 | Wild | Italian wild boar | European wild boar | 9.8 | PRJEB9922 |
| ERS804972 | Wild | Japanese wild boar | Asian wild boar | 9.0 | PRJEB9922 |
| SRS703307 | Wild | Korean wild boar | Asian wild boar | 9.8 | PRJNA260763 |
| SRS703309 | Wild | Korean wild boar | Asian wild boar | 9.4 | PRJNA260763 |
| SRS703310 | Wild | Korean wild boar | Asian wild boar | 10.0 | PRJNA260763 |
| SRS703311 | Wild | Korean wild boar | Asian wild boar | 9.1 | PRJNA260763 |
| SRS703312 | Wild | Korean wild boar | Asian wild boar | 8.5 | PRJNA260763 |
| SRS703313 | Wild | Korean wild boar | Asian wild boar | 6.8 | PRJNA260763 |
| SRS703314 | Wild | Korean wild boar | Asian wild boar | 9.0 | PRJNA260763 |
| SRS703316 | Wild | Korean wild boar | Asian wild boar | 8.1 | PRJNA260763 |
| SRS703308 | Wild | Korean wild boar | Asian wild boar | 10.6 | PRJNA260763 |
| SRS703315 | Wild | Korean wild boar | Asian wild boar | 9.1 | PRJNA260763 |
| ERS177345 | Wild | Netherlandish wild boar | European wild boar | 6.5 | PRJEB1683 |
| ERS177346 | Wild | Netherlandish wild boar | European wild boar | 6.7 | PRJEB1683 |
| ERS177348 | Wild | Netherlandish wild boar | European wild boar | 5.4 | PRJEB1683 |
| ERS805013 | Wild | Netherlandish wild boar | European wild boar | 9.5 | PRJEB9922 |
| ERS805015 | Wild | Netherlandish wild boar | European wild boar | 8.0 | PRJEB9922 |
| ERS805016 | Wild | Netherlandish wild boar | European wild boar | 10.9 | PRJEB9922 |
| ERS805017 | Wild | Netherlandish wild boar | European wild boar | 9.5 | PRJEB9922 |
| ERS805018 | Wild | Netherlandish wild boar | European wild boar | 13.9 | PRJEB9922 |
| ERS805019 | Wild | Netherlandish wild boar | European wild boar | 5.5 | PRJEB9922 |
| ERS805020 | Wild | Netherlandish wild boar | European wild boar | 6.7 | PRJEB9922 |
| ERS805021 | Wild | Netherlandish wild boar | European wild boar | 8.4 | PRJEB9922 |
| ERS805022 | Wild | Netherlandish wild boar | European wild boar | 8.0 | PRJEB9922 |
| ERS805023 | Wild | Netherlandish wild boar | European wild boar | 12.2 | PRJEB9922 |
| ERS805024 | Wild | Netherlandish wild boar | European wild boar | 10.9 | PRJEB9922 |
| ERS177350 | Wild | Swiss wild boar | European wild boar | 5.7 | PRJEB1683 |
| ERS805039 | Wild | Ukrainian Wild | European wild boar | 7.4 | PRJEB9922 |

**Table S7.**

**Sample details of domestic and wild rabbits in this study.** The Run ID and sample name were obtained from the Metadata file in the NCBI SRA Run Selector.

| **Run ID** | **Sample name** | **Wild/Domestic** | **Group Fig. S8** | **Total reads** | **T**  **(reference)** | **Deletion**  **(ancestral)** | **C** | **Insertion*T** |
| --- | --- | --- | --- | --- | --- | --- | --- | --- |
| SRR1290760 | Domestic_FrenchLop_1 | Domestic | French Lop | 8 | 7 | 0 | 0 | 1 |
| SRR1290783 | Domestic_Dutch_1 | Domestic | Dutch | 9 | 1 | 8 | 0 | 0 |
| SRR1290816 | Domestic_Champagne_1 | Domestic | Champagne | 10 | 9 | 1 | 0 | 0 |
| SRR1290820 | Domestic_NewZealand_1 | Domestic | New Zealand White | 8 | 8 | 0 | 0 | 0 |
| SRR1290821 | Domestic_FlemishGiant_1 | Domestic | Flemish Giant | 20 | 19 | 0 | 0 | 1 |
| SRR1290823 | Domestic_BelgianHare_1 | Domestic | Belgian Hare | 11 | 9 | 1 | 0 | 1 |
| SRR1290763 | WildIberian_M_1 | Wild | Iberian | 10 | 2 | 8 | 0 | 0 |
| SRR1290766 | WildIberian_Castanar_1 | Wild | Iberian | 13 | 2 | 11 | 0 | 0 |
| SRR1290768 | WildIberian_CO_1 | Wild | Iberian | 9 | 0 | 9 | 0 | 0 |
| SRR1290782 | WildIberian_TO_1 | Wild | Iberian | 17 | 5 | 12 | 0 | 0 |
| SRR1290810 | WildIberian_Calzada_1 | Wild | Iberian | 12 | 3 | 9 | 0 | 0 |
| SRR1290811 | WildIberian_Guadalajara_1 | Wild | Iberian | 14 | 0 | 13 | 1 | 0 |
| SRR1290812 | WildFrench_Caumont | Wild | French | 14 | 4 | 10 | 0 | 0 |
| SRR1290813 | WildIberian_SCMora_1 | Wild | Iberian | 15 | 6 | 9 | 0 | 0 |
| SRR1290815 | WildFrench_Villemolaque_1 | Wild | French | 6 | 3 | 3 | 0 | 0 |
| SRR1290817 | WildIberian_Toledo_1 | Wild | Iberian | 6 | 0 | 6 | 0 | 0 |
| SRR1290819 | WildFrench_LaRoque_1 | Wild | French | 17 | 3 | 14 | 0 | 0 |
| SRR1290822 | WildIberian_Huelva | Wild | Iberian | 9 | 1 | 8 | 0 | 0 |
| SRR1290825 | WildIberian_Carrion_1 | Wild | Iberian | 13 | 3 | 10 | 0 | 0 |
| SRR1290826 | WildIberian_Zaragoza_1 | Wild | Iberian | 6 | 1 | 5 | 0 | 0 |

**Table S8.**

**The domestic and wild sheep used in this study.** Sequencing depth is given relative to the sheep reference genome.

| **Run ID** | **Domestic/Wild** | **Group Fig. S9** | **Depth** | **BioProject** |
| --- | --- | --- | --- | --- |
| ERR4413943 | Domestic | Merino | 12.4 | PRJEB39179 |
| SRR3938071 | Domestic | Merino | 2.8 | PRJNA325682 |
| SRR3938072 | Domestic | Merino | 3.8 | PRJNA325682 |
| SRR3938073 | Domestic | Merino | 3.6 | PRJNA325682 |
| SRR3938074 | Domestic | Merino | 3.6 | PRJNA325682 |
| SRR5991186 | Domestic | Merino | 4.6 | PRJNA325682 |
| SRR5991236 | Domestic | Merino | 5.0 | PRJNA325682 |
| SRR5991254 | Domestic | Merino | 4.8 | PRJNA325682 |
| SRR3938079 | Domestic | Merino | 3.0 | PRJNA325682 |
| SRR3938095 | Domestic | Merino | 4.6 | PRJNA325682 |
| SRR3938107 | Domestic | Merino | 3.2 | PRJNA325682 |
| SRR3938116 | Domestic | Merino | 2.2 | PRJNA325682 |
| SRR3938128 | Domestic | Merino | 4.4 | PRJNA325682 |
| SRR3938152 | Domestic | Merino | 4.2 | PRJNA325682 |
| SRR5991409 | Domestic | Merino | 2.8 | PRJNA325682 |
| SRR5991185 | Domestic | Merino | 11.8 | PRJNA325682 |
| SRR5991184 | Domestic | Merino | 14.0 | PRJNA325682 |
| SRR5991187 | Domestic | Merino | 12.8 | PRJNA325682 |
| SRR5991272 | Domestic | Merino | 11.6 | PRJNA325682 |
| SRR5991165 | Domestic | Merino | 13.2 | PRJNA325682 |
| SRR5991167 | Domestic | Merino | 13.0 | PRJNA325682 |
| SRR5991170 | Domestic | Merino | 11.4 | PRJNA325682 |
| SRR5991171 | Domestic | Merino | 11.0 | PRJNA325682 |
| SRR5991327 | Domestic | Merino | 8.2 | PRJNA325682 |
| SRR5991326 | Domestic | Merino | 12.8 | PRJNA325682 |
| SRR5991323 | Domestic | Merino | 13.8 | PRJNA325682 |
| SRR5991322 | Domestic | Merino | 14.0 | PRJNA325682 |
| SRR5991155 | Domestic | Merino | 8.6 | PRJNA325682 |
| SRR5991372 | Domestic | Merino | 13.2 | PRJNA325682 |
| SRR5991191 | Domestic | Merino | 10.4 | PRJNA325682 |
| SRR5991273 | Domestic | Merino | 11.4 | PRJNA325682 |
| SRR5991475 | Domestic | Merino | 10.4 | PRJNA325682 |
| SRR5991157 | Domestic | Merino | 10.0 | PRJNA325682 |
| SRR5991168 | Domestic | Merino | 9.4 | PRJNA325682 |
| SRR5991172 | Domestic | Merino | 7.2 | PRJNA325682 |
| SRR5991173 | Domestic | Merino | 6.6 | PRJNA325682 |
| SRR5991325 | Domestic | Merino | 8.2 | PRJNA325682 |
| SRR5991324 | Domestic | Merino | 12.4 | PRJNA325682 |
| SRR5991321 | Domestic | Merino | 12.8 | PRJNA325682 |
| SRR5991166 | Domestic | Merino | 8.2 | PRJNA325682 |
| SRR5991174 | Domestic | Merino | 13.8 | PRJNA325682 |
| SRR5991330 | Domestic | Merino | 8.6 | PRJNA325682 |
| SRR5991156 | Domestic | Merino | 12.2 | PRJNA325682 |
| SRR5991158 | Domestic | Merino | 7.6 | PRJNA325682 |
| SRR5991229 | Domestic | Merino | 10.6 | PRJNA325682 |
| SRR5991248 | Domestic | Merino | 11.8 | PRJNA325682 |
| SRR5991249 | Domestic | Merino | 10.8 | PRJNA325682 |
| SRR5991256 | Domestic | Merino | 11.0 | PRJNA325682 |
| SRR5991370 | Domestic | Merino | 11.8 | PRJNA325682 |
| SRR5991472 | Domestic | Merino | 11.0 | PRJNA325682 |
| SRR5991190 | Domestic | Merino | 10.8 | PRJNA325682 |
| SRR5991363 | Domestic | Merino | 8.8 | PRJNA325682 |
| SRR5991188 | Domestic | Merino | 10.0 | PRJNA325682 |
| SRR5991252 | Domestic | Merino | 7.4 | PRJNA325682 |
| SRR5991366 | Domestic | Merino | 8.0 | PRJNA325682 |
| SRR5991250 | Domestic | Merino | 12.6 | PRJNA325682 |
| SRR5991195 | Domestic | Merino | 8.2 | PRJNA325682 |
| SRR5991255 | Domestic | Merino | 13.6 | PRJNA325682 |
| SRR5991247 | Domestic | Merino | 11.4 | PRJNA325682 |
| SRR5991160 | Domestic | Merino | 12.4 | PRJNA325682 |
| SRR5991365 | Domestic | Merino | 14.4 | PRJNA325682 |
| SRR5991364 | Domestic | Merino | 14.0 | PRJNA325682 |
| SRR5991368 | Domestic | Merino | 12.8 | PRJNA325682 |
| SRR5991367 | Domestic | Merino | 9.2 | PRJNA325682 |
| SRR5991480 | Domestic | Merino | 10.0 | PRJNA325682 |
| SRR5991477 | Domestic | Merino | 14.2 | PRJNA325682 |
| SRR5991474 | Domestic | Merino | 9.2 | PRJNA325682 |
| SRR5991473 | Domestic | Merino | 12.2 | PRJNA325682 |
| SRR5991476 | Domestic | Merino | 9.2 | PRJNA325682 |
| SRR5991253 | Domestic | Merino | 8.4 | PRJNA325682 |
| SRR5991251 | Domestic | Merino | 7.6 | PRJNA325682 |
| SRR5991369 | Domestic | Merino | 11.0 | PRJNA325682 |
| SRR5991371 | Domestic | Merino | 10.8 | PRJNA325682 |
| SRR5991481 | Domestic | Merino | 10.8 | PRJNA325682 |
| SRR5991478 | Domestic | Merino | 11.0 | PRJNA325682 |
| SRR5991479 | Domestic | Merino | 10.6 | PRJNA325682 |
| SRR5991189 | Domestic | Merino | 10.0 | PRJNA325682 |
| SRR5991159 | Domestic | Merino | 14.0 | PRJNA325682 |
| SRR5991500 | Domestic | Merino | 12.4 | PRJNA325682 |
| SRR5991332 | Domestic | Merino | 11.6 | PRJNA325682 |
| SRR5991331 | Domestic | Merino | 13.4 | PRJNA325682 |
| SRR5991496 | Domestic | Merino | 13.8 | PRJNA325682 |
| SRR5991408 | Domestic | Merino | 13.2 | PRJNA325682 |
| SRR5991164 | Domestic | Merino | 13.4 | PRJNA325682 |
| SRR5991461 | Domestic | Merino | 10.0 | PRJNA325682 |
| SRR5991231 | Domestic | Merino | 11.2 | PRJNA325682 |
| SRR5991505 | Domestic | Merino | 11.0 | PRJNA325682 |
| SRR5991507 | Domestic | Merino | 11.4 | PRJNA325682 |
| SRR5991407 | Domestic | Merino | 11.4 | PRJNA325682 |
| SRR5991404 | Domestic | Merino | 9.4 | PRJNA325682 |
| SRR5991402 | Domestic | Merino | 10.6 | PRJNA325682 |
| SRR5991506 | Domestic | Merino | 11.8 | PRJNA325682 |
| SRR5991504 | Domestic | Merino | 12.8 | PRJNA325682 |
| SRR5991314 | Domestic | Merino | 5.8 | PRJNA325682 |
| SRR5991464 | Domestic | Merino | 10.0 | PRJNA325682 |
| SRR5991163 | Domestic | Merino | 14.0 | PRJNA325682 |
| SRR5991162 | Domestic | Merino | 12.6 | PRJNA325682 |
| SRR5991169 | Domestic | Merino | 13.8 | PRJNA325682 |
| SRR5991345 | Domestic | Merino | 9.2 | PRJNA325682 |
| SRR5991458 | Domestic | Merino | 11.8 | PRJNA325682 |
| SRR5991406 | Domestic | Merino | 12.4 | PRJNA325682 |
| SRR5991344 | Domestic | Merino | 14.0 | PRJNA325682 |
| SRR5991405 | Domestic | Merino | 8.0 | PRJNA325682 |
| SRR5991460 | Domestic | Merino | 13.2 | PRJNA325682 |
| SRR5991161 | Domestic | Merino | 8.0 | PRJNA325682 |
| SRR5991462 | Domestic | Merino | 10.2 | PRJNA325682 |
| SRR5991463 | Domestic | Merino | 14.6 | PRJNA325682 |
| SRR5991350 | Domestic | Merino | 13.6 | PRJNA325682 |
| SRR5991349 | Domestic | Merino | 14.2 | PRJNA325682 |
| SRR5991347 | Domestic | Merino | 11.2 | PRJNA325682 |
| SRR5991352 | Domestic | Merino | 14.0 | PRJNA325682 |
| SRR5991351 | Domestic | Merino | 14.4 | PRJNA325682 |
| SRR5991233 | Domestic | Merino | 13.8 | PRJNA325682 |
| SRR5991175 | Domestic | Merino | 13.8 | PRJNA325682 |
| SRR5991459 | Domestic | Merino | 11.4 | PRJNA325682 |
| SRR5991456 | Domestic | Merino | 10.8 | PRJNA325682 |
| SRR5991457 | Domestic | Merino | 10.0 | PRJNA325682 |
| SRR5991465 | Domestic | Merino | 11.0 | PRJNA325682 |
| SRR5991348 | Domestic | Merino | 11.8 | PRJNA325682 |
| SRR5991346 | Domestic | Merino | 10.4 | PRJNA325682 |
| SRR5991343 | Domestic | Merino | 11.6 | PRJNA325682 |
| SRR5991228 | Domestic | Merino | 11.2 | PRJNA325682 |
| SRR5991199 | Domestic | Merino | 11.8 | PRJNA325682 |
| SRR5991230 | Domestic | Merino | 10.2 | PRJNA325682 |
| SRR5991232 | Domestic | Merino | 11.6 | PRJNA325682 |
| SRR5991234 | Domestic | Merino | 10.6 | PRJNA325682 |
| SRR5991235 | Domestic | Merino | 11.2 | PRJNA325682 |
| SRR12396950 | Wild | Urial | 19.9 | PRJNA645671 |
| CRR053800 | Wild | European mouflon | 4.0 | PRJCA001227 |
| SRR12396880 | Wild | Argali | 15.6 | PRJNA645671 |
| SRR12396881 | Wild | Argali | 17.6 | PRJNA645671 |
| SRR12396882 | Wild | Argali | 20.2 | PRJNA645671 |
| SRR12396883 | Wild | Argali | 17.3 | PRJNA645671 |
| SRR12396885 | Wild | Argali | 16.0 | PRJNA645671 |
| SRR12396886 | Wild | Argali | 15.7 | PRJNA645671 |
| SRR12396887 | Wild | Argali | 16.1 | PRJNA645671 |
| SRR12396951 | Wild | Argali | 18.2 | PRJNA645671 |
| CRR053545 | Wild | Argali | 5.3 | PRJCA001221 |
| CRR053546 | Wild | Argali | 4.4 | PRJCA001221 |
| CRR053801 | Wild | Argali | 3.8 | PRJCA001227 |
| SRR11657500 | Wild | Asian mouflon | 28.6 | PRJNA624020 |
| SRR11657501 | Wild | Asian mouflon | 25.0 | PRJNA624021 |
| SRR11657502 | Wild | Asian mouflon | 24.1 | PRJNA624022 |
| SRR11657503 | Wild | Asian mouflon | 23.4 | PRJNA624023 |
| SRR11657504 | Wild | Asian mouflon | 33.6 | PRJNA624024 |
| SRR11657505 | Wild | Asian mouflon | 25.9 | PRJNA624025 |
| SRR11657506 | Wild | Asian mouflon | 22.1 | PRJNA624026 |
| SRR11657507 | Wild | Asian mouflon | 22.8 | PRJNA624027 |
| SRR11657635 | Wild | Asian mouflon | 27.9 | PRJNA624028 |
| SRR11657636 | Wild | Asian mouflon | 22.9 | PRJNA624029 |
| SRR11657637 | Wild | Asian mouflon | 23.4 | PRJNA624030 |
| SRR11657638 | Wild | Asian mouflon | 26.4 | PRJNA624031 |
| SRR11657639 | Wild | Asian mouflon | 25.0 | PRJNA624032 |
| SRR11657640 | Wild | Asian mouflon | 32.9 | PRJNA624033 |
| SRR11657641 | Wild | Asian mouflon | 22.4 | PRJNA624034 |
| SRR11657642 | Wild | Asian mouflon | 23.6 | PRJNA624035 |
| ERR157930 | Wild | Asian mouflon | 13.3 | PRJEB3139 |
| ERR157931 | Wild | Asian mouflon | 13.0 | PRJEB3140 |
| ERR157932 | Wild | Asian mouflon | 12.6 | PRJEB3141 |
| ERR157938 | Wild | Asian mouflon | 13.3 | PRJEB3142 |
| ERR157939 | Wild | Asian mouflon | 11.9 | PRJEB3143 |
| ERR157942 | Wild | Asian mouflon | 13.0 | PRJEB3144 |
| ERR157944 | Wild | Asian mouflon | 13.1 | PRJEB3145 |
| ERR332573 | Wild | Asian mouflon | 14.8 | PRJEB3146 |
| ERR332575 | Wild | Asian mouflon | 13.8 | PRJEB3147 |
| ERR332582 | Wild | Asian mouflon | 14.8 | PRJEB3148 |
| ERR332589 | Wild | Asian mouflon | 13.1 | PRJEB3149 |
| ERR340346 | Wild | Asian mouflon | 12.5 | PRJEB3150 |
| ERR466544 | Wild | Asian mouflon | 12.1 | PRJEB3151 |
| ERR157935 | Wild | Asian mouflon | 12.8 | PRJEB3152 |
| ERR315509 | Wild | Asian mouflon | 12.6 | PRJEB3153 |
| ERR332587 | Wild | Asian mouflon | 12.6 | PRJEB3154 |
| ERR466546 | Wild | Asian mouflon | 12.6 | PRJEB3155 |

**Table S9.**

**Frequency distribution of the pLoPD mutation rs317817652*T in all five subspecies of red junglefowl.** Data were obtained from Galbase (<http://animal.omics.pro/code/index.php/ChickenVar>).

| Subspecies | Number | Frequency(rs317817652*T) |
| --- | --- | --- |
| *Gallus gallus bankiva* | 4 | 0 |
| *Gallus gallus gallus* | 6 | 0 |
| *Gallus gallus jabouillei* | 27 | 0 |
| *Gallus gallus murghi* | 60 | 0 |
| *Gallus gallus spadiceus* | 44 | 0.193 |

**Table S11.**

**pLoPD mutations in goats, horses, and dogs.** The data were obtained from the Ensembl database. Asterisks denote that these variants have been reported to be associated with body size and are present in Fig. 3.

| Species | Genome | PALI2 protein | pLoPD variant | Consequence of pLoPD variant |
| --- | --- | --- | --- | --- |
| Horse | Equcab3.0 | ENSECAP00000056810.1 | *rs1146838995 | ENSECAP00000056810.1:p.Glu818Ter |
| Goat | ARS1 | ENSCHIP00000021571.1 | *rs657074013 | ENSCHIP00000021571.1:p.Ser277IlefsTer38 |
|  |  |  | rs658083174 | ENSCHIP00000021571.1:p.Ala663ArgfsTer17 |
|  |  |  | rs644167651 | ENSCHIP00000021571.1:p.Arg773Ter |
|  |  |  | rs638969232 | ENSCHIP00000021571.1:p.Tyr1060Ter |
|  |  |  | rs645991632 | ENSCHIP00000021571.1:p.Asn1180IlefsTer13 |
|  |  |  | rs664155883 | ENSCHIP00000021571.1:p.Leu1395TyrfsTer11 |
| Dog | ROS_Cfam_1.0 | ENSCAFP00845000412.1 | *rs3327936124 | ENSCAFP00845000412.1:p.Arg1257LysfsTer12 |
|  |  |  | rs3327944765 | ENSCAFP00845000412.1:p.Pro478CysfsTer6 |
|  |  |  | rs3327944699 | ENSCAFP00845000412.1:p.Pro478SerfsTer6 |
|  |  |  | rs3327779413 | ENSCAFP00845000412.1:p.Asp661AsnfsTer22 |

Table S1. Information for 3,208 CLUES-identified selected SNVs.

Table S2. 158 cattle body size QTLs used for co-localization with CLUES-identified selected SNVs.

Table S3. Information for 111 CLUES-identified selected SNVs co-localized with 11 body size QTLs.

Table S4. Information for SNVS and INDELs in A and D haplotypes.

Table S5. Information for the 10 variants with the largest frequency differences between Haplotypes A and D.

Table S10. rs317817652*T significantly increases body weight and body size in Chinese local chickens.

Table S12. *LCORL* and *STC2* mutations frequencies in different breeds of *Bos taurus* and *Bos indicus*. Only *Bos taurus* breeds with a sample size greater than 10 were included in the analysis.
